# Metacells untangle large and complex single-cell transcriptome networks

**DOI:** 10.1101/2021.06.07.447430

**Authors:** Mariia Bilous, Loc Tran, Chiara Cianciaruso, Aurélie Gabriel, Hugo Michel, Santiago J. Carmona, Mikael J. Pittet, David Gfeller

## Abstract

The exponential scaling of scRNA-seq data represents an important hurdle for downstream analyses. Here we develop a coarse-graining framework where highly similar cells are merged into metacells. We demonstrate that metacells not only preserve but often improve the results of downstream analyses including visualization, clustering, differential expression, cell type annotation, gene correlation, imputation, RNA velocity and data integration. By capitalizing on the redundancy inherent to scRNA-seq data, metacells significantly facilitate and accelerate the construction and interpretation of single-cell atlases, as demonstrated by the integration of 1.46 million cells from COVID-19 patients in less than two hours on a standard desktop.

## Background

Single-cell RNA sequencing (scRNA-seq) provides transcriptome-wide gene expression profiles at single-cell resolution. This technology has been transformative for unsupervised investigation of heterogeneous cell populations [1,2], identification of novel cell states and cell types [3], discovery of novel markers [4] and reconstruction of developmental lineages [5].

The rapid developments of single-cell capture and sequencing technologies enable researchers to profile tens to hundreds of thousands of single cells [6–9]. This high throughput provides unprecedented opportunities to explore the heterogeneity of complex tissues [6,10]. For instance, even very rare cell types can be detected by unsupervised clustering [11–13]. However, the large number of cells that need to be profiled from biological tissues in order to capture their complete heterogeneity is an important hurdle for downstream analyses.

Several approaches to streamline the analysis of large scRNA-seq data have focused on improving bioinformatics pipelines to scale with more cells [14–17] and adapt them to computational infrastructures that can cope with large memory requirements [18,19]. These developments often require the use of dedicated platforms, which are less user-friendly and not necessarily available to researchers without in-depth training in bioinformatics. As an alternative, cell subsampling approaches have been developed [20,21]. However, subsampling does not capitalize on the full information present in the initial scRNA-seq data, which is likely sub-optimal, for instance to reduce the noise inherent to scRNA-seq data due to dropout [22].

High-throughput single-cell transcriptomic profiling of biological samples typically leads to the repetitive sampling of highly similar, and possibly biologically redundant cells. Some attempts to simplify scRNA-seq data by merging such highly similar cells into metacells have been proposed, including the MetaCell algorithm [23,24]. MetaCell has been successfully used for visualization and exploratory purposes [25–27], but does not scale well with very large numbers of cells. Moreover, it remains unclear whether metacells can be used for quantitative and robust downstream analyses, and how much biological information may be gained or lost when analyzing such simplified data [28].

Here, we developed a network-based coarse-graining framework to simplify scRNA-seq data by merging highly similar cells into metacells. We demonstrate that metacells (i) preserve the global structure of the initial data, (ii) enable efficient and robust downstream analyses, as demonstrated by the identification of genes specifically expressed in tumor-infiltrating dendritic cell subtypes, (iii) significantly reduce noise from single-cell gene expression measurements and (iv) lead to ten- to hundred-fold reduction of the size of the data and the computational time and memory requirements.

## Results

### Simplifying scRNA-seq data with metacells

To facilitate the analysis of scRNA-seq data, we developed a computational coarse-graining framework, called SuperCell, based on the idea of grouping highly similar cells into metacells (Fig. 1a). First, scRNA-seq data are modeled as a single-cell k-nearest neighbor (kNN) graph with nodes representing cells and edges connecting cells with high transcriptomic similarity [29,30] (see Methods). Next, metacells are built by merging single cells with very high internal connectivity. To this end, we used the walktrap algorithm [31], which allows users to predefine the number of metacells, although other algorithms may be used. Unlike standard clustering, our aim is not to identify cell populations that can be mapped to distinct biological cell types, but rather to merge cells that contain highly similar and likely repetitive transcriptomic information. The graining level (*γ*) is defined as the ratio between the number of cells and the number of metacells. Finally, a metacell gene expression matrix is computed by averaging gene expression within metacells (Fig. 1a).

**Figure 1.**
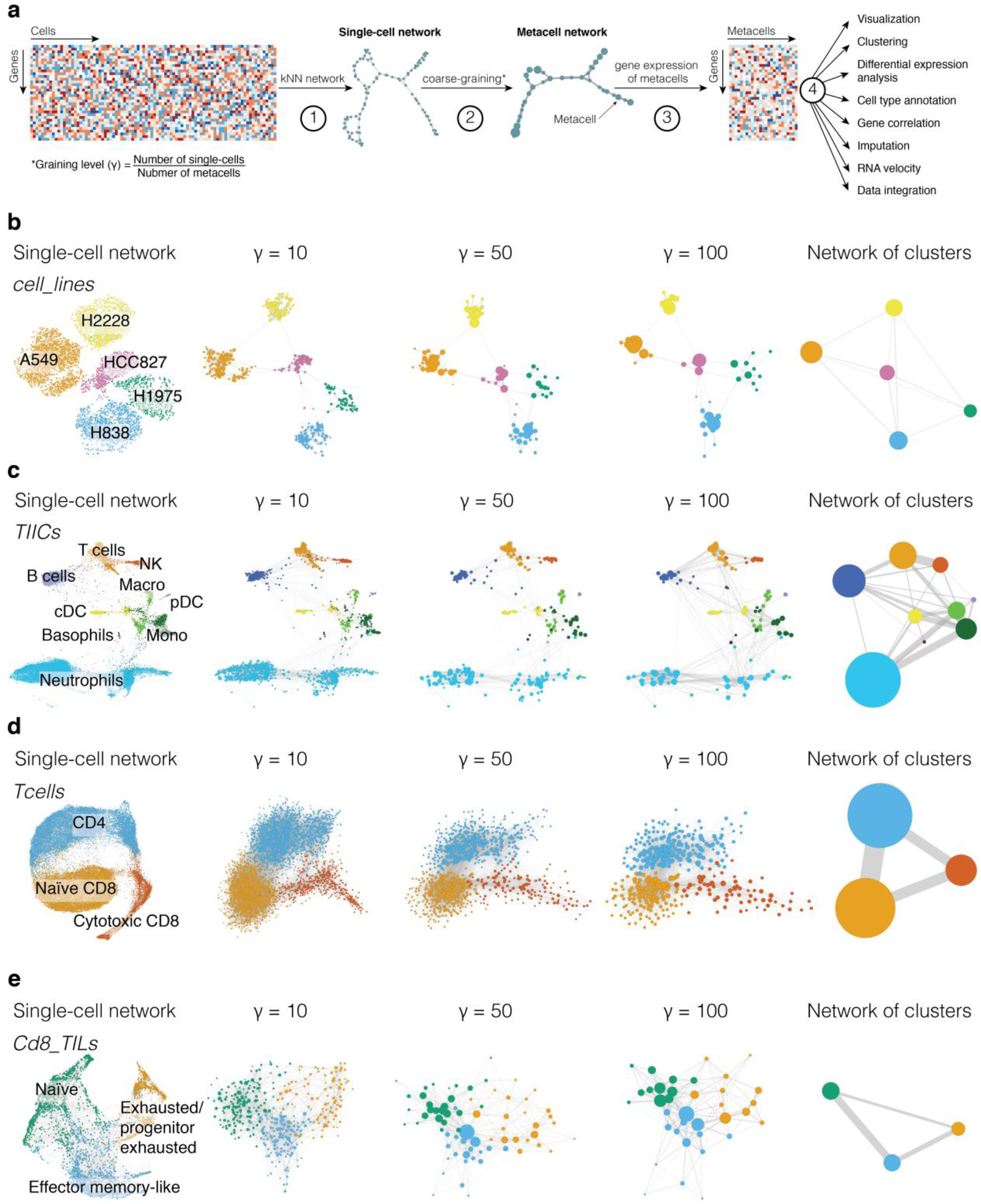
Simplifying single-cell RNA-seq data with SuperCell. **a**, Overview of the SuperCell coarse-graining pipeline, including the following steps. (1) A single-cell network is constructed from the single-cell gene expression matrix using k-nearest neighbors (kNN) algorithm. (2) Densely connected cells are merged into metacells at a user-defined graining level (*γ*). (3) A gene expression matrix of metacells is computed by averaging gene expression within each metacell. (4) The metacell gene expression matrix can be used for visualization and downstream analyses such as clustering, differential expression, cell type annotation, gene correlation, imputation, RNA velocity and data integration. **b**-**e**, Examples of metacell networks at several graining levels. For comparison, the network of clusters is shown on the right. (**b**) Five cancer cell lines (*cell_lines, N* = 3’918) shown with different colors. (**c**) Tumor-infiltrating immune cells (*TIICs, N* = 15′939). (**d**) T cells sorted from PBMC (*Tcells, N* = 40’560). (**e**) Tumor-infiltrating CD8 T lymphocytes (*Cd8_TILs, N* = 3’574).

To explore and benchmark the SuperCell algorithm, we first applied it to four scRNA-seq datasets and compared the results of the analysis at the single-cell and metacell levels. The first dataset consists of five human adenocarcinoma cell lines (*cell_lines, N* = 3′918) [32] (Fig. 1b). The second one consists of tumor-infiltrating immune cells from murine KP1.9 lung adenocarcinoma (*TIICs, N* = 15′939) [4] (Fig. 1c). The third one consists of purified T cells from healthy donors (*Tcells, N* = 40′560) [9] (Fig. 1d). The fourth one consists of tumor-infiltrating CD8 T lymphocytes (*Cd8_TILs, N* = 3′574) [33] (Fig. 1e). The *cell_lines* dataset represents a gold standard for clustering where the ground truth corresponds to the different cell lines. The other three datasets represent more biologically relevant cases, spanning different levels of heterogeneity from CD45^+^ cells, to T cells to subsets of CD8 T cells. Colors in Fig. 1b-c correspond to the cell type annotations retrieved in the original studies. Colors in Fig. 1d-e represent clusters in single cells since cell type annotation was not unambiguously known. In Fig. 1d (*Tcells*), the three clusters could be mapped to CD4, naïve CD8 and cytotoxic CD8 T cells. In Fig. 1e (*Cd8_TILs*), the three clusters correspond to naïve, effector memory-like, and exhausted/progenitor-exhausted CD8 T cells.

The metacells at different graining levels preserve the global structure of the single-cell data and are compatible with different types of visualization techniques based on networks (Fig. 1b-e) or dimensionality reduction (Supplementary Fig. 1). As expected, the size of metacells increases with higher graining levels (Supplementary Fig. 2a). The average fraction of genes detected in each metacell also increases with *γ*, indicating that metacells are less prone to the high dropout typically observed in single cells (Supplementary Fig. 2b).

To further assess how much metacells preserve the structure of the single-cell data, we checked whether metacells contain cells originating from the same cell type. To this end, we used the purity, defined as the proportion of the most abundant cell type in a metacell (see Method). Fig. 2a shows that metacells have purity close to 1, indicating that they consist mainly of cells of the same cell type. This high purity allowed us to annotate metacells according to the most abundant cell type in each metacell (colors in Fig. 1b-e). As a negative control (i.e., lower bound), random grouping of cells would result in much lower purity (Fig. 2a). We further compared our results to those obtained with MetaCell [23]. We used either the default mode (MetaCell_def) or a SuperCell-like mode (MetaCell_SC) which uses the same set of genes and the same way of averaging gene expression as in SuperCell (see Methods). The purity obtained with both versions of MetaCell is either equivalent or lower than with SuperCell (Fig. 2a).

**Figure 2.**
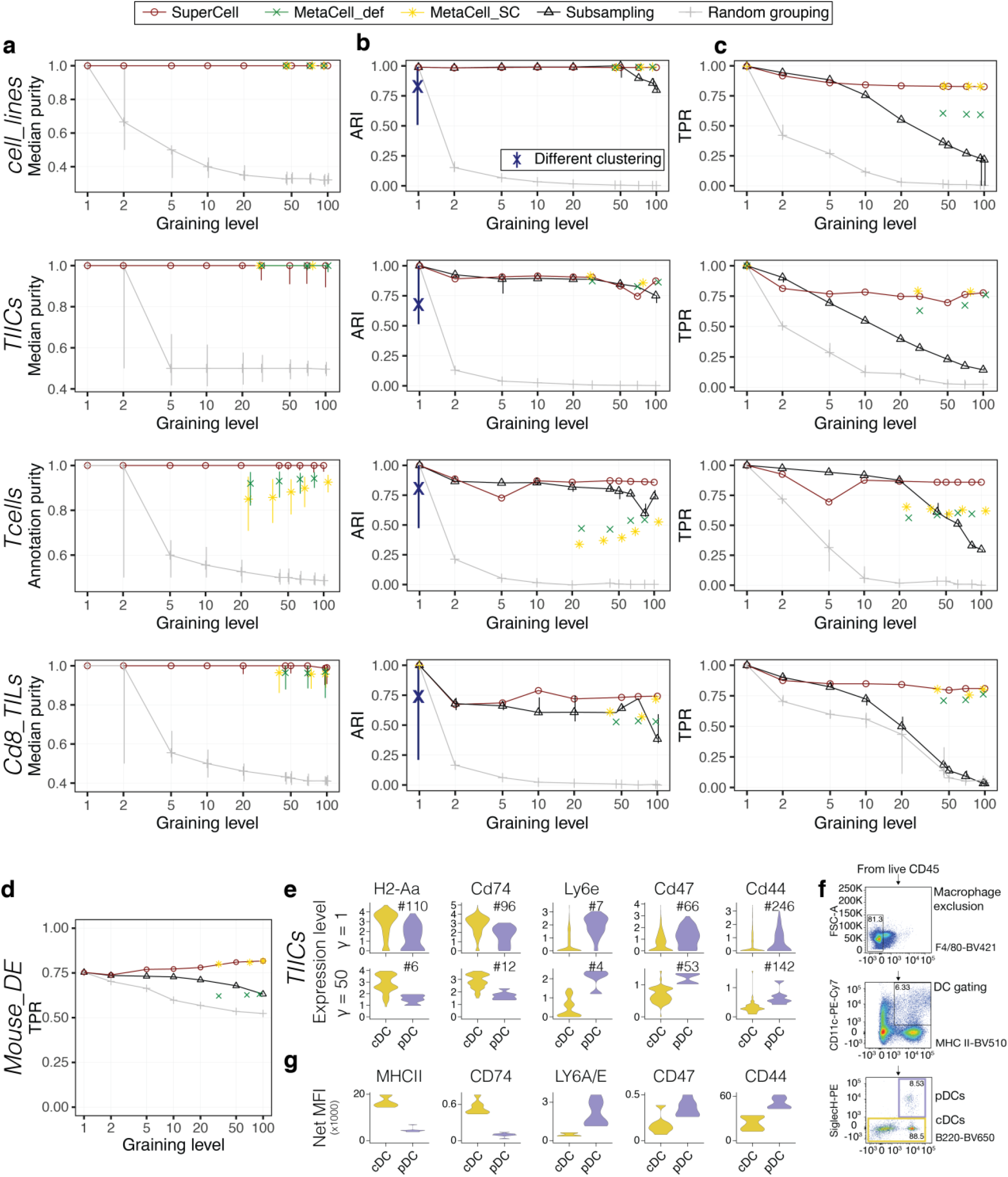
Metacells preserve clustering and differential expression results, and reveal genes specifically expressed in dendritic cell subtypes. **a**, Median purity of metacells computed with SuperCell, MetaCell_def and MetaCell_SC at different graining levels for the four datasets shown in Fig. 1b-e (*cell_lines*, *TIICs*, *Tcells*, *Cd8_TILs*). As a lower bound, the purity after random grouping of cells is shown in gray. **b**, Consistency between the hierarchical clustering of metacells or after subsampling and the one of single cells (see Supplementary Fig. 4a for results with other clustering algorithms). The blue line shows the range of ARI values when other clustering algorithms are applied to the single-cell data (median shown with “X”). **c**, Proportion of the cluster-specific DE genes (based on weighted t-test) found at the single-cell level and recovered at the metacell level or after subsampling. **d**, Proportion of the condition-specific DE genes found in bulk RNA-seq and recovered at the metacell level or after subsampling in the *Mouse_DE* dataset. **e**, Expression of genes coding for trans-membrane proteins in single cells (top) and metacells (bottom) that are more differentially expressed (i.e., better ranking) between cDCs and pDCs at the metacell level. The number following the ‘#’ sign indicates the ranking of each gene among the top differentially expressed ones. **f**, Flow cytometry analysis of DCs from murine KP1.9 lung adenocarcinoma (*N* = 7). **g**, Median fluorescence intensity of proteins coded by the genes from **e**. All comparison shown in **e** and **g** pass statistical significance based on two-tailed unpaired Student’s t-test (p-values < 0.05).

To confirm that the overall high purity also applies to rare cell types, we took advantage of the *TIICs* dataset which contains two rare cell populations: plasmacytoid dendritic cells (representing < 0.5% of the cells) and basophils (representing < 0.2% of the cells). We observed that both the proportion of single cells from these rare cell types that are mapped to metacells of the same cell type, as well as the purity of these metacells are high across multiple graining levels (see Methods and Supplementary Fig. 3).

### Metacells preserve clustering results

Beyond visualization, an important step in scRNA-seq data analysis is to identify distinct cell types or cell states by clustering. To check the consistency of the clustering of metacells with the clustering of single cells, we used the adjusted Rand index (ARI) (see Methods). Our results shows that ARI values are high across different granularity levels and better than subsampling for large *γ* (Fig. 2b). This applies especially to the gold standard *cell_lines* dataset where single-cell clusters represent each a distinct cell line. To further investigate the consistency between the clustering at the single-cell and metacell levels, we compared it to the consistency of clusters obtained when using different algorithms on the single-cell data. We observed that ARI values for clustering of the metacells are within the same range as ARI values for the clustering of single cells using different clustering algorithms (Fig. 2b, blue vertical line). These results indicate that metacells can be used for clustering, and the differences with the clustering at the single-cell level are within the expected fluctuations observed with different choices of clustering algorithms. ARI values were similar or higher for metacells built with SuperCell compared to those built with MetaCell. Similar results were obtained with other clustering algorithms (Supplementary Fig. 4a).

We then explored whether the original number of clusters, defined as the one that maximizes the silhouette coefficient [34], can be recovered in metacells. Supplementary Fig. 4b demonstrates that this number could in general be recapitulated and this conservation compares favorably with subsampling. Additionally, even in cases where the predicted optimal number of clusters differs between single cells and metacells, the clusters of metacells are still consistent with the ones of single cells (Supplementary Fig. 4c).

As metacells contain different numbers of single cells, we used a sample-weighted hierarchical clustering algorithm in Fig. 2b. To explore the impact of weights on metacells, we compared the performance of unweighted versus sample-weighted clustering. Overall, Supplementary Fig. 5 shows that both unweighted and sample-weighted clustering algorithms perform similarly.

We finally explored other choices of parameters and methods in the SuperCell pipeline (see Methods). Altogether, we observed lower performance with higher values of k, when using shared nearest neighbor (sNN) instead of kNN networks or when using the Louvain clustering algorithm instead of the walktrap (Supplementary Fig. 6). These observations suggest that the default parameters in SuperCell provide a reasonable solution to build metacells, although we do not exclude that other solutions could lead to similar performance.

### Metacells are consistent with differential expression analysis

Another use of scRNA-seq data analysis is to identify differentially expressed genes between clusters or conditions. To explore the performance of differential expression (DE) analysis between clusters at the metacell level for the four datasets of Fig. 1b-e, we used sample-weighted t-test and assessed the recovery rate of differentially expressed genes found at the single-cell level using the true positive rate (TPR) (see Methods). Our results show that more than 75% of the DE genes observed at the single-cell level can be recovered in metacells even at relatively high graining levels (Fig. 2c), in contrast to subsampling or random grouping. Improvements were also observed with SuperCell compared to MetaCell. Comparison of sample-weighted versus unweighted DE revealed similar TPR for moderate granularity (*γ* < 50), while for larger *γ*, we observed improved performance when using sample-weighted t-test (Supplementary Fig. 7). As with clustering consistency, other choices of parameters and methods in the SuperCell pipeline did not improve the results (Supplementary Fig. 8).

We next explored the results of DE between different conditions. To this end, we capitalized on a dataset (*Mouse_DE*) generated to benchmark DE between conditions (treated vs untreated) in scRNA-seq data compared to bulk RNA-seq [35]. We assessed the recovery of DE genes found in the bulk at both the single-cell and metacell level, using EdgeR [36]. We observed that metacells built either with SuperCell or MetaCell improved recovery of DE genes found in bulk (Fig. 2d). Similar results were obtained with other DE algorithms, like DESeq2 [37] or t-test, or when using AUC instead of TPR (Supplementary Fig. 9). This demonstrates that metacells can significantly reduce the size of the data and simultaneously identify DE genes between conditions that better recapitulate those found in bulk.

### Metacells reveal genes specifically expressed in dendritic cell subtypes

To further illustrate the use of metacells for the identification of genes expressed in specific cell types, we examined the *TIICs* dataset (Fig. 1c). We performed DE analysis between conventional (cDCs) and plasmacytoid (pDCs) dendritic cells, which are known to play important roles in eliciting the immune response against cancer, at both single-cell and metacell levels. Several genes displayed clearer DE patterns in metacells (i.e., better ranking). These include *H2-Aa*, which is known to be upregulated in cDCs [38,39] (ranked #6 in the metacell DE analysis, while it was ranked #110 in the single-cell DE analysis, Fig. 2e). To experimentally validate these predictions, we selected additional genes coding for trans-membrane proteins with available antibodies and that were ranked better in the metacell DE analysis (see Methods). These included *Cd74* (upregulated in cDCs) as well as *Ly6e, Cd47* and *Cd44* (upregulated in pDCs) (Fig. 2e). We then performed a flow cytometry analysis of DCs of the same murine lung cancer model (Fig. 2f and Supplementary Fig. 10, see Methods). The results confirmed the DE of all the selected genes at the protein level (Fig. 2g), demonstrating that metacells are useful for magnifying biological information in scRNA-seq data that is less detectable at the single-cell level.

### Metacells improve cell type annotation, gene correlation and imputation

We next compared marker-based cell type annotation at single-cell and metacell levels. For this, we first annotated CD4 and CD8 T cells from the *Tcells* dataset using either single markers (Fig. 3a) or gene signatures derived from bulk RNA-seq data (Fig. 3b, see Methods) and compared this annotation with the cell type determined by protein expression during the sorting procedure (see Methods). The annotation quality, computed as the area under the ROC curve (AUC), grows with the graining level until it almost reaches saturation (AUC=1) for both CD4 and CD8 T cells. Annotations based on metacells obtained with MetaCell lead to lower AUC. As expected, subsampling failed to improve cell type annotation. Similar results were obtained when annotating metacells in the *Cd8_TILs* dataset (Supplementary Fig. 11). Inaccuracies in annotating cells using single markers or gene signatures are likely due to the high dropout rate [22]. In the *Tcells* dataset, less than 40% of CD4, respectively CD8 T cells, express the *CD4*, respectively *CD8A*, marker genes (Fig. 3c). By contrast, at the graining level *γ* = 100, 79%of CD4 metacells and 97%of CD8 metacells express *CD4* and *CD8A* markers, respectively.

**Figure 3.**
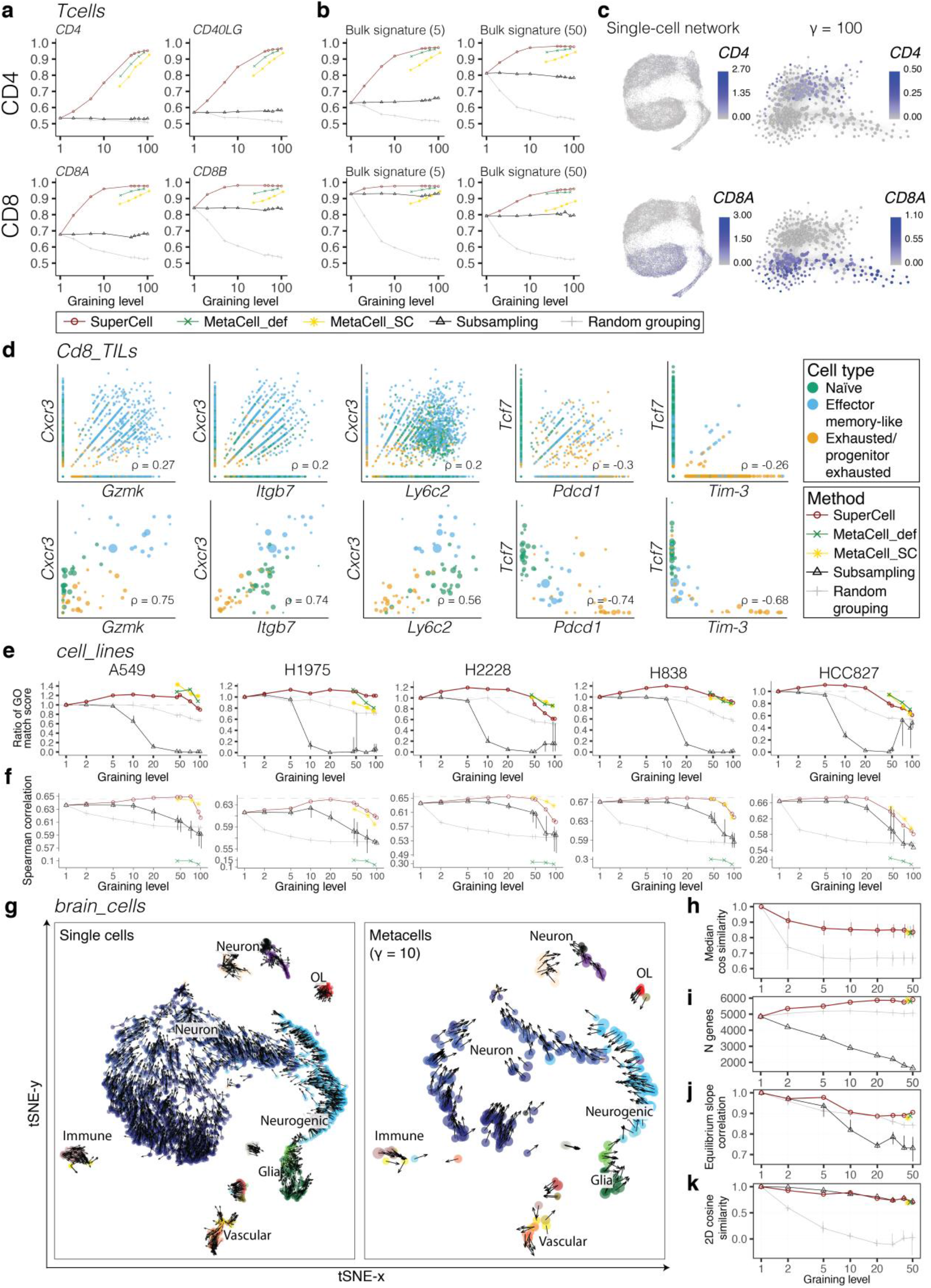
Metacells improve cell type annotation, gene correlation, imputation and RNA velocity. **a**-**b**, AUC of recovery of CD4 (top) and CD8 (bottom) T cells from the *Tcells* dataset using single markers (**a**) or signatures defined form bulk (**b**) consisting of the top 5 or top 50 genes for metacells computed with SuperCell, MetaCell_def, MetaCell_SC and random grouping, or after subsampling. **c**, Expression of *CD4* (top) and *CD8A* (bottom) in T cells from the *Tcells* dataset at the single-cell and the metacell (*γ* = 100) levels. **d**, Gene correlation at the single-cell (top) and the metacell (*γ* = 50) (bottom) levels for selected gene pairs in the *Cd8_TILs* dataset, with the corresponding sample-weighted Pearson correlation (ρ). **e**, Comparison of the GO similarity of metacell and single-cell top correlated genes identified for individual cell lines from the *cell_lines* dataset. The y-axis shows the ratio between mean GO match scores of the top correlated genes at the metacell and the single-cell levels. **f**, Mean Spearman correlation between bulk and MAGIC-imputed data in each cell line of the *cell_lines* dataset. The dashed lines show the correlation between the pseudo-bulk (i.e., averaged gene expression within a cell line) and bulk gene expression. **g**, Joint tSNE visualization of RNA velocity for the *brain_cells* dataset (N = 3’396) for single cells (left) and metacells (*γ* = 10) (right) colored by cell type annotation. **h**, Velocity purity in metacells (defined as the cosine similarity of single-cell velocities within each metacell). **i**, Number of genes with valid estimated equilibrium slope values. **j**, Pearson correlation of gene equilibrium slope values obtained in single-cell and metacell RNA velocities. **k**, Cosine similarity between 2D single-cell and metacell RNA velocities shown in **g**. For the subsampling and random grouping, the center of the error bars denotes the median, and the extrema denotes the 1^st^ and 3^rd^ quartiles (obtained with different random seeds).

We next investigated gene correlation, which is an important source of information to infer gene co-expression modules [40]. We first used the *Cd8_TILs* dataset and explored genes that are known to be positively correlated (e.g., *Ibgb7, Cxcr3, Ly6c*, and *Gzmk* which are markers of effector memory-like cells) or negatively correlated (e.g., *Tcf7* that is expressed in naïve and stem-like cells, versus *Pdcd1* and *Havcr2/Tim-3* that are expressed in exhausted cells). Fig. 3d shows that metacells increase expected gene correlations and remove the noise arising from single-gene dropouts in scRNA-seq. To more systematically explore the biological relevance of correlated genes, we compared the biological relatedness of top correlated gene pairs found exclusively at the single-cell or metacell levels within individual cell lines in the *cell_lines* dataset. As a measure of similarity between a pair of genes, we used a Gene Ontology (GO) [41] match score (see Methods). Fig. 3e shows higher values of the average GO match score of the top correlated gene pairs in the metacells compared to the single cells until a graining level of roughly 50. Average GO match score computed in metacells built with Supercell or MetaCell had similar behavior, although the limited range of graining levels in MetaCell prevented comparisons for *γ* values where the largest improvement was observed with SuperCell. Overall, these results show that pairs of correlated genes found in metacells show higher biological relatedness compared to those found in single cells.

Imputation methods were shown to improve signal-to-noise ratio in scRNA-seq data and lead to better correlation with bulk profiles [42]. To investigate whether metacells can be used as an input for imputation approaches, we applied MAGIC [40] to both the single cells and the metacells from the *cell_lines* dataset. The results show that the imputed gene expression profiles are more similar to the bulk profiles when applying MAGIC on metacells (Fig. 3f). The best improvement is reached within the same range of graining levels as for gene-gene correlations (Fig. 3e). The results obtained with MetaCell_SC are comparable to those of SuperCell, while the correlations obtained with MetaCell_def are much lower. Overall, our results indicate that imputation can be applied to metacells and leads to improved correspondence with bulk data.

### Metacells are compatible with RNA velocity

We next investigated whether metacells can be used to study differentiation processes with RNA velocity [5]. We first considered a dataset of mouse hippocampus cells (*brain_cells, N* = 3′396) [43] and applied the velocyto algorithm [5] to single cells and metacells (see Methods). We observed consistent RNA velocity results at the metacell (right) and the single-cell (left) levels (Fig. 3g) when plotted on a joint t-distributed stochastic neighbor embedding (tSNE). Both cases show the developmental path from neurogenic cells (light blue) to neurons (dark blue). This consistency is further confirmed with a high purity of metacells in terms of velocity across multiple graining levels (see Methods) (Fig. 3h). To compute RNA velocity, a key parameter is the estimated equilibrium slope of each gene (i.e., linear fit between spliced and un-spliced mRNA, referred to as “γ” in the original RNA velocity publication [5]). The number of genes for which this equilibrium slope can be estimated increases with higher graining levels for metacells, while it stays constant for random grouping, and decreases for subsampling (Fig. 3i). Moreover, the equilibrium slopes of genes in metacells correlate with those in single cells, and the correlation is higher than for the subsampling (Fig. 3j). The improvements of metacells over subsampling are likely due to the regularizing and enriching effects of metacells on spliced and un-spliced mRNA abundance (see examples in Supplementary Fig. 12a). To directly compare RNA velocity at the single-cell and metacell levels in the tSNE plots of Fig. 3g, we computed the cosine similarity between 2D RNA velocity of each single cell and 2D RNA velocity of the metacell it belongs to (Fig. 3k). This similarity score also suggests a high consistency of metacell RNA velocity across multiple graining levels. In the joint tSNE used in Fig. 3g to facilitate comparison, subsampling displayed similar conservation of the 2D RNA velocity (Fig. 3k). However, when plotted separately, the differentiation process is much more visible in metacells than after subsampling (Supplementary Fig. 12b). Metacells built with MetaCell also preserve the results of RNA-velocity, demonstrating the robustness of the metacell concept for RNA-velocity analysis in scRNA-seq data. The same analyses were performed on a mouse pancreas scRNA-seq dataset (*pancreatic_cells, N* = 3′696) [44] and similar results were obtained (Supplementary Fig. 13). This demonstrates that metacells are compatible with RNA velocity.

### Metacells facilitate data integration

ScRNA-seq atlases are built by integrating data from multiple samples. To explore the use of metacells in scRNA-seq data integration, we analyzed a recently published dataset of 1.46 million immune cells coming from various tissues from 196 COVID-19 patients and healthy controls (*COVID-19_atlas, N* = 1′462′702) [45]. Analyzing these data is challenging since most existing data integration algorithms do not scale with such cell numbers on standard computational infrastructures. We applied the SuperCell algorithm on each sample separately (see Methods), which led to a total of 146’304 metacells (Fig. 4a). We then performed data integration on metacells with Harmony [46] (Fig. 4b). The results of integrated metacells showed high similarity with those obtained based on single cells in the original study. In particular, the main immune cell types reported in the original study could be recapitulated. Visually, Fig. 4b shows that protocols and samples are well mixed after integration of metacells. This is confirmed by the improvement of the kBET acceptance rates (a quantitative measure of batch mixing) [47] both in terms of protocols (Fig. 4c, top) and samples (Fig. 4c, bottom). The clustering of integrated data has also higher consistency with the original cell type annotation (ARI=0.75, compared to ARI=0.66 in non-integrated data). To further illustrate the use of metacells, we performed DE analysis followed by gene set enrichment analysis within monocytes and B cells from COVID-19 patients versus healthy controls, at the metacell level (see Methods). For monocytes, the most upregulated GO term in COVID-19 patients included ‘chemokine mediated signaling pathway’ (GO:0070098, adjusted p-value = 1.3 · 10^−7^) and ‘complement activation’ (GO:0006956, adjusted p-value = 4.1 · 10^−5^). The most upregulated GO terms in B cells from COVID-19 patients included ‘inflammatory response’ (GO:0006954, adjusted p-value = 5.9 · 10^−3^), ‘type I interferon signaling’ (GO:0060337, adjusted p-value = 3.7 · 10^−2^) or ‘response to viruses’ (GO:0009615, adjusted p-value = 3.7 · 10^−2^), consistent with a humoral response to SARS-Cov2. Importantly, all these analyses could be run on a standard desktop (Macbook Pro i7 core, 16G RAM) in less than two hours.

**Figure 4.**
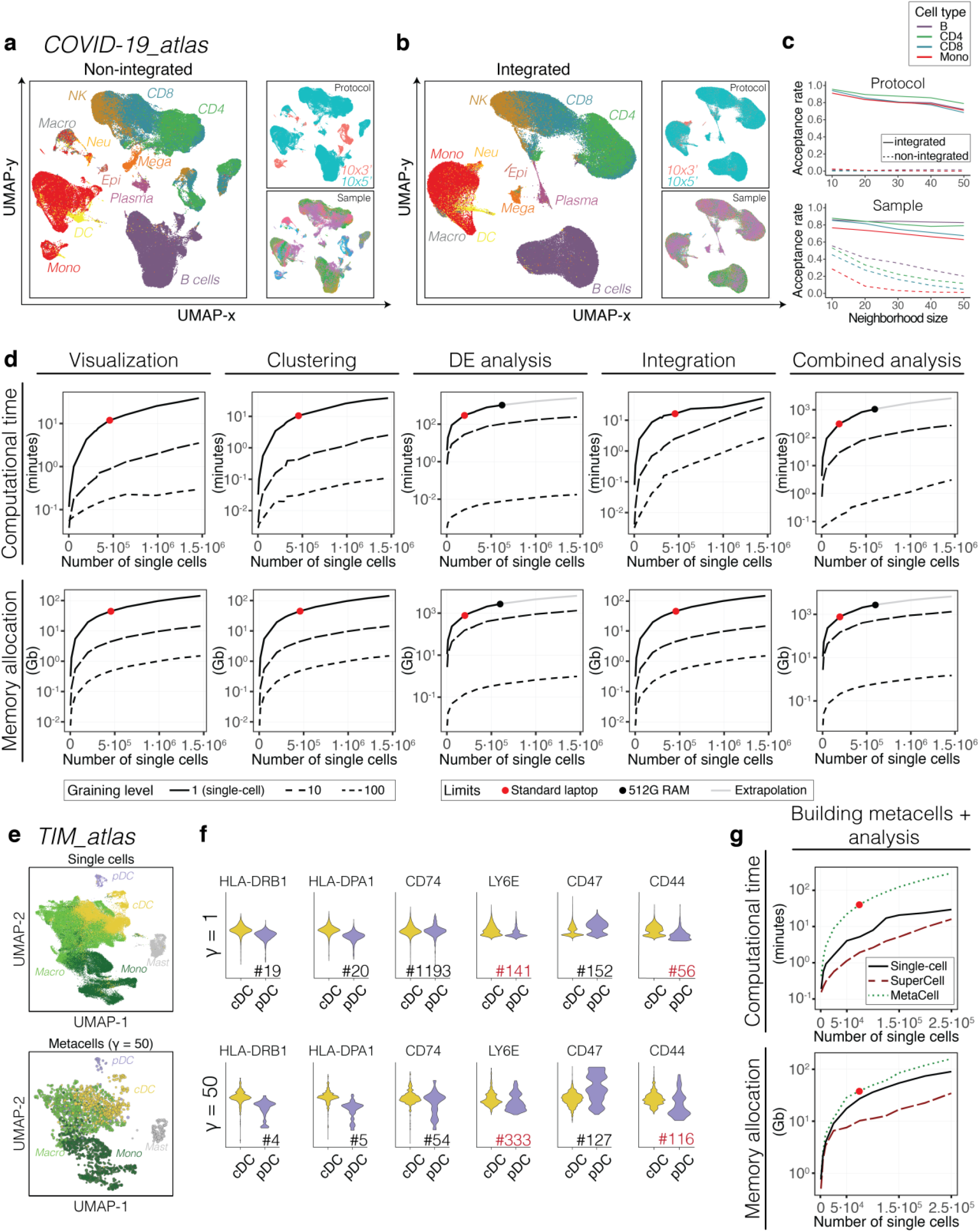
Metacells facilitate data integration and accelerate downstream analyses. **a**-**b**, UMAP visualization of the non-integrated (**a**) and Harmony-integrated (**b**) *COVID-19_atlas* dataset (*N* = 1’462’702) at the metacell (*γ* = 10) level. Metacells are colored according to the cell type annotation, protocol or sample. **c**, Batch effect level in terms of protocol (top) and sample (bottom) in the non-integrated and Harmony-integrated *COVID-19_atlas* dataset, computed as the kBET acceptance rate for the four most frequent cell types. **d**, Computational time (top) and memory allocation (bottom) for the visualization (UMAP), clustering (Seurat), DE analysis (t-test, each cell type versus the rest), data integration (Harmony) and all steps together (‘Combined analysis’) for the metacells (dashed lines) and single cells (solid line). Red dots show the limits reached on standard desktops (16G of RAM). Black dots correspond to the limits reached on a machine with 512G RAM (linear extrapolations shown in gray). **e**, UMAP visualization of the *TIM_atlas* dataset (*N* = 108’566) at the single-cell (left) and the metacell (*γ* = 50) (right) levels computed with the approximate coarse-graining. Cells are colored according to the cell type annotation. **f**, Relative (z-score) expression of genes experimentally tested in Fig. 2g at the single-cell (top) and metacell (bottom) levels. The number following the ‘#’ sign indicates the ranking of each gene among the top differentially expressed ones. All comparisons pass statistical significance based on two-tailed unpaired Student’s t-test (p-values < 0.05) except for *CD74* at the single-cell level (p-value = 1). Ranks for genes showing a different behavior both at single-cell and metacell levels between mouse and human are shown in red. **g**, Computational time (top) and memory allocation (bottom) for the building of metacells followed by downstream analyses including dimensionality reduction, clustering and DE analysis for metacells computed with SuperCell (red dashed line) or MetaCell (green dashed line), and for the single cells (solid black line). Red dots show the limit reached on standard desktops (16G of RAM).

### Metacells significantly accelerate downstream analyses and reduce memory requirements

To assess the improvement in computational efficiency obtained with metacells, we benchmarked the time and memory needed for the downstream analyses, including visualization, clustering, DE analysis (i.e., each major cell type versus the rest) and data integration (Fig. 4d), as well as RNA velocity (Supplementary Fig. 14) (see Methods). As expected, each analysis runs faster on metacells and uses less memory. All analyses could be run on a standard desktop with 16G of RAM at the metacell level, while they crashed at ~200′000 cells for DE and ~480′000 cells for the other analyses at the single-cell level (red dots in Fig. 4d). Moreover, DE analysis could not be performed at the single-cell level for datasets with more than 600′000 cells even on a high-performance computing (HPC) platform with 512G of RAM (black dots in Fig. 4d, extrapolations in gray). This demonstrates the advantage of metacells for exploratory analysis of large scRNA-seq datasets.

Although metacell construction needs to be run only once, it can be computationally demanding for large numbers of cells (> 100’000). To address this issue, we included an option to perform an approximate coarse-graining in SuperCell, which first builds metacells using a subset of cells and then maps the rest of the cells to the most similar metacells (Supplementary Fig. 15a) (see Methods). Supplementary Fig. 15b-e show that the approximate version of SuperCell has similar performance as the exact one. To further demonstrate the ability of the approximate coarse-graining to deal with large-scale scRNA-seq datasets, we applied it to a human pan-cancer atlas of tumor-infiltrating myeloid cells (*TIM_atlas, N* = 108′566) [39] (Fig. 4e). Focusing on DCs, we performed DE analysis between cDCs and pDCs for the genes tested in Fig. 2g. Our results confirm the improved DE signal in metacells compared to single cells for *HLA-DRB1, HLA-DPA1, CD74* and *CD47* (Fig. 4f). The signal for *LY6E* and *CD44* was opposite to their expression pattern in mouse at both the single-cell and metacell levels, suggesting that their expression in DCs subtypes may not be conserved between human and mouse or across cancer types.

Taking everything together, the entire analysis, including building of metacells, dimensionality reduction, clustering and DE analysis, runs faster with metacells than with single cells and requires less memory (Fig. 4g, Supplementary Fig. 16a,b). Compared to MetaCell, our approach is significantly faster and can be applied to larger datasets, both when considering the entire analysis (Fig. 4g), or only the construction of metacells (Supplementary Fig. 16c). These results indicate that metacells significantly facilitate and accelerate the analysis of large scRNA-seq data.

## Discussion

ScRNA-seq technologies are revolutionizing biological sciences by providing transcriptome-wide information for very large numbers of individual cells. Here we introduce the SuperCell coarse-graining pipeline for scRNA-seq data. We demonstrate that metacells built with SuperCell preserve the properties of the single-cell data at multiple graining levels and serve as a compromise structure between the single-cell level and the level of clusters. The fact that metacells are compatible with clustering, differential expression, cell type annotation, gene correlation, imputation, RNA velocity and data integration indicates that this framework can be readily applied to the majority of analyses performed on scRNA-seq data. Moreover, metacells can magnify biologically relevant information, as demonstrated with the identification and validation of markers of tumor-infiltrating dendritic cell subtypes (Fig. 2e-g). When exploring scRNA-seq data, different methods or choices of parameters are typically tested for visualization, clustering, differential expression, RNA velocity or data integration [48,49]. The SuperCell framework is therefore especially appropriate for such exploratory analyses.

Unlike clustering, the primary aim of metacells is not to identify groups of cells with a specific biological interpretation (e.g., cell types or cell states), but to simplify, accelerate and improve the analysis of scRNA-seq data. As such, the exact number of metacells is not meant to have a specific biological significance and can be fixed by the users based on the available computational resources. For practical applications, we recommend using *γ* ∈ [10,50], as this already provides a significant speed-up in the analysis of large datasets and preserves downstream analyses results. We also note that defining optimal *γ* with measures like the silhouette coefficient used in clustering would result in very large values where properties of the scRNA-seq data are no longer conserved.

For the construction of metacells, we used the walktrap algorithm. Owing to its hierarchical structure, this algorithm enables users to explore different graining levels without having to recompute the metacells for each choice of *γ*. This can be useful considering the heterogeneity in size and complexity of scRNA-seq datasets. Comparison with the Louvain clustering algorithm suggests that the walktrap algorithm provides a reasonable solution (Supplementary Fig. 6 and 8). However, it is expected that robust metacells can be built with other approaches since the exact definition of each metacell is not critical, as long as cells of high transcriptomic similarity are merged, and the results of downstream analyses are preserved.

The high purity of metacells indicates that they mainly consist of cells of the same cell type. However, we also observed that a small fraction cells from rare cell types can be mixed with cells from other cell types in some metacells. To overcome this issue, we implemented the possibility to build metacells in a way that is consistent with *a priori* defined cell type annotations (see Methods). We anticipate that this option will significantly facilitate the re-analysis of large datasets carefully annotated based on expert knowledge which is difficult to perfectly recapitulate with unsupervised analyses.

The metacell concept shares similarity with other computational approaches developed for scRNA-seq data analysis. Akin to imputation [40], it averages signals over cells with high transcriptomic similarity. However, results of imputations can be difficult to use with very large datasets, since the total number of cells remains the same and the imputed gene expression matrices are less sparse than the original ones. Methods analyzing networks of clusters, like PAGA [30] or TooManyCells [50], can be thought of as a coarse-graining with a very high graining level. These approaches have been developed for other purposes and by construction cannot preserve the intra-cluster heterogeneity. Large values of the graining level (e.g., in order of *N*/10) also lead to significant loss of information in downstream analyses, as well as technical challenges (e.g., DE between clusters cannot be robustly performed if each cluster is made of one single metacell). Compared to the MetaCell algorithm [51], where the concept of metacells was first introduced, the SuperCell approach displays equal or better performance, is more flexible in terms of graining levels, runs faster and scales better with larger numbers of cells.

Beyond the details of the metacell construction procedure, our work provides a robust framework for systematic benchmarking of the results obtained with metacells. This is critical to ensure that metacells can indeed be used without losing important information from the single-cell data. As such, we believe that this framework will be useful to benchmark other coarse-graining approaches that will be or are being developed [52,53].

## Conclusions

Altogether, our work shows that SuperCell significantly reduces the size of scRNA-seq data and accelerates downstream analyses, while preserving the global structure of the data and the results of such downstream analyses. These results demonstrate that metacells provide an intermediate and tunable level of representation for scRNA-seq data between the single-cell level, which is partly redundant and suffers from high technical noise, and the cluster level, which can mask biologically relevant intracluster heterogeneity. As the throughput of single-cell technologies keeps increasing [54], we anticipate that SuperCell-like approaches will play an increasingly important role in facilitating visualization, analysis, sharing and interpretation of single-cell genomics data.

## Methods

### Metacell construction

Metacells are built based on a log-normalized gene expression matrix. The set of features (genes) used for the construction of metacells is defined by default as the set of the most variable genes but can also be provided by the user. Based on this set of features, a low dimensional embedding is computed using principal component analysis (PCA) (function *irlba*() from *irlba* R package [55]). The top principal components (top 10 by default in this work) are used to build a single-cell network with a k-nearest neighbors (k-NN, *k* = 5 for all studied datasets) algorithm (*nn2()* function from *RANN* R package [56]): each cell (node) is connected to *k* most similar cells based on the Euclidean distance. To construct metacells, we apply the *walktrap* clustering algorithm (available as *clust_walktrap()* function from *igraph* R package [57]) to the singe-cell network. Metacells are constructed by merging single cell (nodes) at a user-specified graining level (*γ*). The graining level is defined as the ratio between the number of single cells (*N_c_*) and the number of metacells (*N_SC_*). In the metacell network, the size of a metacell is defined as the total number of cells it contains. The weight of an edge connecting two metacells is computed as the total number of edges connecting cells of those metacells. A gene expression profile of metacells is computed by averaging gene expression within metacells.

The SuperCell method is implemented in an R package (SuperCell) that can be used for all steps of the downstream analyses and is compatible with other state-of-the-art tools for scRNA-seq data analyses, including Seurat [15] and SingleCellExperiment [14].

### Purity of metacells

The purity of a metacell is defined as the proportion of cells from the most abundant cell type in this metacell. For the *cell_lines* and *TIICs* datasets, we used the cell type annotation provided in the original studies [4]. For the *Tcells* and *Cd8_TILs* datasets, we annotated single cells based on clustering since cell type annotation is not known a priori. When studying conservation of rare cell types (i.e., plasmacytoid dendritic cells or basophils) in the *TIICs* dataset, metacells were annotated to a particular cell type with the Bayesian classifier developed in Zillionis et al. [4]. We then computed the proportion of single cells of each of the two rare cell types that were found in metacells annotated to the same cell type (Supplementary Fig. 3a). For the subsampling, this value was computed as a proportion of subsampled single cells of the rare cell type of interest. We further computed the purity of each metacell annotated to these rare cell types (Supplementary Fig. 3b). This purity was defined as the fraction of single cells of the corresponding cell type weighted by the number of these single cells to account for heterogeneity in the size of metacells.

### PCA in metacells

A scaling of the metacell gene expression matrix was applied before PCA based on a weighted version of a scale function (*wt.scale()* function from *corpcor* R package [58]). Next, a singular value decomposition (SVD) was computed (function *irlba*() from *irlba* R package [55]) for the sample-weighted gene expression matrix 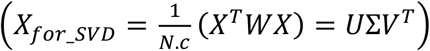, where *X* is a scaled and transposed (cells as rows) gene expression matrix at a metacell level, *W* is a diagonal matrix with sizes of metacells on its diagonal, Σ are the singular vectors of *X_for_SVD_*, *U* and *V* are left- and right-singular vectors of *X_for_SVD_*. Then, the principal component embedding of metacells (*X_PCA_*) was computed as *X_PCA_* = *XV*.

### Clustering consistency

To compute clusters, the top 1000 and 500 variable genes were used to perform the PCA for *cell_lines* and *Tcells* datasets, respectively. For *TIICs* and *Cd8_TILs*, we used the same gene set as in the original studies [4,33]. Single-cell and metacell data were clustered with hierarchical clustering (*hclust()* function in R with the “ward.D” method and the parameter *members* set to the metacell size vector). Alternatively, k-means (*kmeans()* function in R) and Seurat (*FindClusters()* function in *Seurat* package [15]) clustering algorithms were also explored.

To assess the consistency between the clustering of single cells and metacells, Hubert and Arabie’s adjusted Rand index (ARI) was computed. The same number of clusters was used in single cells and metacells. To estimate the variability of ARI values expected with different clustering algorithms at the single-cell level, we clustered single-cell data with alternative clustering algorithms (i.e., different methods of hierarchical clustering, including “ward.D”, “single”, “complete”, “average”, “mcquitty”, “median”, “centroid”, and k-means clustering with 5 different random starts) and computed ARI between the hierarchical clustering and the results obtained with the alternative clustering algorithms (thick blue bars in Fig. 2b).

### Silhouette coefficient in metacells

The computation of the silhouette coefficient was also adjusted for considering the weights of metacells. If *a*(*i*) is the mean distance between a metacell *i* belonging to a cluster *C_m_* and all other metacells from the same cluster, then 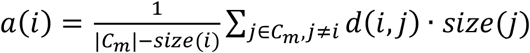, where *d*(*i, j*) is a distance between metacells *i* and *j*, *size*(*i*) is the size of metacell *i*, and |*C_m_*| is the size of cluster *C_m_*. If *b*(*i*) is the minimal average distance from a metacell *i* to any metacell from a different cluster, then 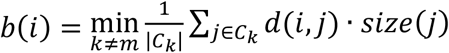. The silhouette of a metacell *i* is 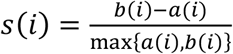 if |*C_m_*| > 1 and *S*(*i*) = 0 if |*C_m_*| = 1. The overall silhouette value *S* is a sample-weighted mean of *s*(*i*), 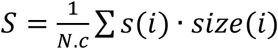.

### Alternatives for metacell construction

To study the sensitivity of the SuperCell algorithm to particular choices of methods and parameters, we built metacells with different values for *k* in kNN single-cell networks, kNN and shared nearest neighbors (sNN) algorithms from Seurat, and Louvain clustering instead of walktrap. Metacells obtained with these different approaches were tested for preservation of the results of clustering (Supplementary Fig. 6) and differential expression (Supplementary Fig. 8).

### Differentially expressed genes recovery rate

To assess the ability of metacells to recover differentially expressed genes, we considered both differential expression between clusters and between conditions. Differential expression between clusters (i.e., each cluster versus all the others) in the *cell_lines*, the *TIICs*, the *Tcells* and the *CD8_TILs* datasets was performed with sample weighted t-test (*wtd.t.test()* function from *weights* R package [59]). Clusters were computed for single cells and metacells. The set of differentially expressed genes in single cells (*M*) was identified as a union of significantly (two-tailed t-test adjusted p-value < 0.05) upregulated (i.e., logFC > 0.5 for *cell_lines* and *Cd8_TILs* or logFC > 0.25 for *TIICs* and *Tcells*) genes from all clusters. The set of differentially expressed genes at the metacell level 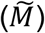 was computed as the top *n* = |*M*| significantly (i.e., two-tailed t-test adjusted p-value < 0.05) upregulated genes across all clusters ranked by logFC.

Differential expression between conditions (i.e., control versus treated samples) in the *Mouse_DE* dataset was performed with t-test, DESeq2 [37] and EdgeR [36]. For each test, the set of the differentially expressed genes (*M*) was computed from the bulk RNA-seq dataset as the set of significantly (adjusted p-value < 0.05) differentially expressed genes. The set of differentially expressed genes at the metacell level 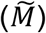 was computed as the top *n* = |*M*| significantly (i.e., adjusted p-value < 0.05) differentially expressed genes ranked by absolute value of logFC.

To measure the conservation of DE genes, the True Positive Rate (TPR) was computed as 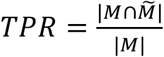. This value is equivalent to the precision since 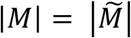. For the differential expression analysis between conditions, the AUC was additionally computed using the adjusted p-value as a predictor and the set of differentially expressed genes in bulk as a label.

### Benchmarking with MetaCell

The MetaCell algorithm [23] was run in two modes: a default (green) and a SuperCell-like (yellow) mode. For the default mode, MetaCell was run using the vignette instructions provided at the authors’ GitHub (https://github.com/tanaylab/metacell/tree/master/vignettes). As a gene expression matrix of MetaCell, we used the output field of MetaCell named *mc_fp* (MetaCell footprint). For the SuperCell-like mode, MetaCell was run using the same set of genes used with the SuperCell approach and the same way of computing the gene expression (i.e., by averaging gene expression within metacells). Since MetaCell does not explicitly allow users to tune the graining level, we had to vary the parameter of the minimal MetaCell size (*min_mc_size*). This parameter was set to {1,10,*20*,30,50}. This explains why the range of graining level values for MetaCell is narrow.

### Identification of cDCs and pDCs marker genes in the *TIICs* dataset

Data pre-processing and cell type annotation in the *TIICs* dataset at the metacell level was performed the same way as the single-cell analyses in the original study [4]. In brief, metacells were built based on the first 60 principal components computed on highly variable genes (i.e., genes with the variance larger than a mode variance). Metacells were annotated to a particular cell type with a Bayesian classifier developed in Zillionis et al. [4] applied to the metacell count matrix. To plot metacells, we used SPRING layout [29] from the original study averaged within each metacell.

For the annotated *TIICs* dataset, DE analysis between cDCs and pDCs was performed at the single-cell and metacell levels. Among the top 150 differentially expressed genes at the metacell level, we selected those that were detected in both subtypes (i.e., in at least 95% of cDC and pDC metacells) and those that were ranked higher (by p-value significance and then by logFC for identical p-values) at the metacell level (*rank_single−cell_* − *rank_super−cell_* ≥ 3). Among them, genes coding for trans-membrane proteins with available antibodies were tested for their protein expression using flow cytometry (see Supplementary Tables 3 and 4 for the full list of genes matching these criteria).

### Mouse tumor model

Murine KP1.9 lung adenocarcinoma tumor cells were cultured in Iscove’s DMEM media (Corning) supplemented with 10% fetal bovine serum (FBS) and 1% penicillin/streptomycin. Cells were injected into 8 weeks old C57BL/6J male mice (Charles River) intravenously (2.5×105 cells in 100 μl PBS) to develop orthotopic tumors in the lung. The tumor cell line was derived from lung tumor nodules of a C57BL/6 KrasLSL-G12D/WT;p53Flox/Flox (KP) mouse and was kindly provided by Dr. Zippelius (University Hospital Basel, Switzerland). Mice were analyzed for tumor phenotypes 4 weeks post-cancer cell injection [4,60]. All animals were housed at the Agora In Vivo Center (AIVC) in Lausanne. Experiments were performed following protocols approved by the Veterinary Authorities of the Canton Vaud according to Swiss law (animal license VD3612).

### Flow cytometry analysis of mouse lung tumors

Single cell suspensions were obtained from lung tumor tissue of C57BL/6J male mice after transcardial PBS perfusion. Small tissue pieces were generated from perfused lungs using scissors and digested in DMEM containing 2.5% FBS, 25% Accutase (Sigma), 0.5 mg/ml collagenase type IV (Worthington Biochemical Corporation), 0.5 mg/ml hyaluronidase (Sigma) and 5 Units/ml DNAse I (Sigma) for 20 min at 37 C while shaking (800 rpm). Digested lung tissue was gently meshed through 70 μm cell strainers using a plunger. After red blood cells lysis (BD Pharm Lyse), single cell suspensions were incubated with Fc blocker (1:100, BioLegend) in staining buffer (PBS with 2% FBS and 1mM EDTA), followed by cell viability staining with Live/Dead Fixable Zombie UV (1:1000, BioLegend) in PBS. Cells were then stained with fluorochrome-conjugated antibodies (see Supplementary Table 5) for 30 min at 4 C in staining buffer, prior to fixation with IC Fixation buffer (Invitrogen) for 30 min. Samples were acquired using an LSRFortessa apparatus (BD Biosciences), and data were analyzed with FlowJo (v10.7.1). Cells were gated based on their size and granularity (FSC-A vs SSC-A), followed by doublet and dead-cell exclusion. Conventional DCs (cDC) and pDCs were defined as CD45+F4/80–CD11c+MHCII+SiglecH– and CD45+F4/80– CD11 c+MHCII+SiglecH+B220+, respectively.

### Cell type annotation with single genes and gene signature recovered from bulk RNA-seq

Cell type annotation in *Tcells* was performed with either a single marker or a gene signature recovered from bulk RNA (see below). A score was computed as the sum of max-normalized expression of signature genes. AUC was computed using the score as a predictor and CD4/CD8 sorting information as a label with *prediction()* function from *ROCR* R package [61].

To compute CD4/CD8 gene signature from bulk RNA-seq, a dataset of bulk RNA-seq of sorted immune cells from donors were downloaded from GEO under accession number GSE60424 [62]. DE analysis of 20 sorted CD4 and 20 sorted CD8 T cells populations from 10 donors was performed using *edgeR* R package [36]. The disease status was used as a covariate to eliminate disease bias. Genes with *FDR* < 0.01 and |*logFC*| > 0.5 were considered to be true markers (*N* = 550). Only 174 of those markers were found among genes in the *Tcells* dataset. The top 5 and top 50 markers were used as a gene signature of CD4 and CD8 T cell types.

The cell type annotation in *Cd8_TILs* (Supplementary Fig. 11) was performed similarly to the one in *Tcells*, but instead of using the bulk signature, a set of top marker genes of each cluster found at the single-cell level was used. Due to the lack of sorting information in this dataset, the label (ground truth cell type annotation) was set to the single-cell clustering results.

### GO match score of top correlated genes

For each cell line from the *cell_lines* dataset, genes that were expressed in more than 50% of cells of this cell line were kept. The gene pairwise Pearson correlation was computed using a gene expression matrix at the single-cell and at the metacell levels. Then, the top 1′000 statistically significant (adjusted p-value < 0.05) and highly correlated gene pairs at each level were selected. For each gene pair, a GO match score was computed as a Jaccard coefficient of their GO id sets. We then computed an average GO match score for gene pairs found exclusively in metacells and normalized it by an average GO match score of gene pairs found exclusively at the single-cell level (*γ* = 1).

### Imputation in metacells

The imputation in single-cell and metacell data were performed using MAGIC [40] with the default configuration. Bulk RNA-seq data of human adenocarcinoma used for comparison with the imputed profiles were extracted from GEO under accession number GSE86337. Only the common genes between the bulk data and the single cell data (*cell_lines*) were kept for the imputation. For each cell line, the imputed profiles of the metacells and single cells were compared with the corresponding bulk profile using the rank-based Spearman correlation. For the comparison, a pseudo-bulk profile is computed as an average scRNA-seq gene expression within a particular cell line.

### RNA velocity in metacells

Metacells were built in the same way described in the metacell construction section starting from the total counts (sum of spliced and un-spliced counts). The spliced and un-spliced counts for metacells were computed by taking their mean value within each metacell. The RNA velocity of the single cells and metacells was computed independently using *relative.velocity.estimate()* function from the *Velocyto.R* package [5] with fit.quantile = 0.02.

The purity of metacells in terms of velocity was computed based on the cosine similarity of RNA velocities of single cells belonging to one metacell. The purity is determined as a maximum of a median of the cosine between RNA velocity vectors of cells based on the formula:

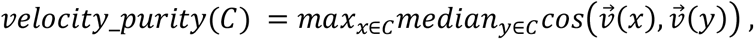

where 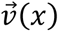 is a velocity of a cell *x* belonging to a metacell *C*.

To obtain comparable 2D velocities vectors between single cells and metacells, a joint tSNE of single-cell and metacell velocities was computed using *tSNE.velocity.plot()* function with the parameters nPCs = 10 and perplexity = 50. We then computed the velocity similarity score as a median of the cosine similarities between the velocity of each single cell and the velocity of corresponding metacell, i.e.,

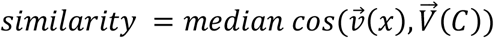

where 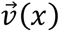 is the velocity of cell *x*, and 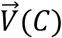 is the velocity of a supercell *C* (that contains *x*) in the joint tSNE. We computed the similarity between single-cell RNA velocity and that of random grouping in the same way. For subsampling method, to make a reasonable comparison, we define the similarity as follows:

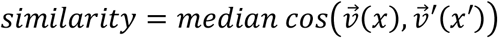

where 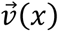 is the velocity of cell *x* and 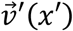 is the velocity of subsampled cell *x*′ that is nearest to *x* (based on the Euclidean distance).

A key parameter estimated with RNA velocity is the equilibrium slope of each gene, which corresponds to the ratio between spliced and un-spliced mRNA that represents the equilibrium between mRNA production and degradation. This equilibrium slope can be computed for a limited number of genes (Fig. 3i). To compare the consistency of RNA velocity results in metacells and single cells, the Pearson correlation between equilibrium slope values was computed for the genes for which this equilibrium slope can be computed in all the methods (metacell, subsampling and random grouping) across all graining levels.

### Data integration of the *COVID-19_atlas* dataset

For each sample of the *COVID-19_atlas* dataset, metacells were first constructed by specifying the graining levels to ensure more than 20 metacells are obtained in each sample:

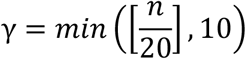

where [] denotes the integer part and *n* denotes the sample size. We then used the *SelectIntegrationFeatures()* function from *Seurat* package [15] to get 2’000 genes for the integration. We next merged the metacell samples into a single object of 146’304 metacells and performed a PCA reduction and Harmony integration [46] of all the samples.

To evaluate the batch effects in terms of protocols and samples before and after integration, kBET acceptance rate [47] was computed for non-integrated and integrated datasets for the cell types present in all samples in a sufficient number (i.e., B cells, monocytes, CD4 T cells and CD8 T cells). UMAP coordinates were used for this estimation.

To compute kBET acceptance rates for protocols, we randomly subsampled *m* metacells from 10×5’ protocol where *m* is number of metacells in 10×3’ protocol. We then applied the function *kBET()* on those subsampled batches for *k* ∈ {10, 20, 30, 40, 50} to obtain the rejection rate *r*(*k*). The kBET acceptance rate is then determined as 1 − *r*(*k*).

The kBET acceptance rates for samples were computed in the same way. In the subsampling step, 15 metacells from each sample were subsampled where possible.

DE analysis within either monocytes or B cells from COVID-19 patients versus healthy controls was performed at the metacell level with Seurat using only PBMC samples. Gene set enrichment was applied to the top 20 differentially expressed genes.

### Approximate coarse-graining for large datasets

For the datasets with size > 100′000, we offer the option to use an approximate coarse-graining in SuperCell, which repeats all the steps of the metacells construction but uses a set of subsampled cells, whose number can be specified by the user, to build an initial subsampled metacell structure. Then, the remaining cells are mapped to the most similar metacell based on the Euclidean distance (Supplementary Fig. 15a).

### Integration of the human *TIM_atlas* dataset

Publicly available and well annotated datasets generated or reported in ref [39] and used to build the atlas of tumor infiltrating myeloid cells (*TIM_atlas, N* = 108′566) are listed in Supplementary Table 2. Integration of the listed datasets was performed as described in the original study [39]. Briefly, we performed two rounds of integration using scanorama package [63]. In the first round, 10x datasets were integrated and corrected. In the second round, corrected 10x data were integrated with all the other datasets.

All the downstream analyses, including metacell construction, cell type annotation and the DE analysis were performed on the corrected gene expression data. Cell type annotation is available for the dataset generated in the original study [39]. Cells from the remaining datasets integrated into the atlas were annotated using a logistic regression, trained on the first 100 principal components of annotated data. Metacells were annotated based on the most abundant cell type in each metacell. The DE analysis between pDC and cDCs was performed at the single-cell and metacell levels the same way as in the *TIICs* dataset.

### Computational time and memory allocation

Assessment of the computational time and memory allocation for the data integration pipeline was performed on the *COVID-19_atlas* dataset. We benchmarked each step of the analyses separately, iteratively increasing the number of single cells by adding new samples. The benchmarking of the visualization includes dimensionality reduction (PCA) and UMAP. The benchmarking of the clustering includes PCA, graph construction and Seurat clustering. The benchmarking of the DE analysis consists of Seurat DE analysis with the t-test for 12 clusters (defined as the main cell types annotated in the original study) and 2′000 genes. Since DE analysis could not be performed for datasets with more than 600′000 cells, the computational time and memory allocation were extrapolated using linear model (gray lines in Fig. 4d). The benchmarking of the data integration consists of PCA and Harmony integration. The benchmarking of the ‘Combined analysis’ consists of all the steps, including PCA, UMAP, graph construction, clustering, DE analysis and Harmony integration. The following functions of the Seurat package [15] were used: *RunPCA(), RunUMAP()*, *RunHarmony()*, *FindNeighbors()* and *FindClusters()*, *FindAllMarkers()*. The benchmarking was performed on one node of an HPC cluster with 512G of RAM. To test the limits reached on standard desktop, the benchmarking was performed on an Intel core i7 with 16G of RAM. To assess computational time and memory allocation, the function *mark()* from R package *bench* [64] was used.

To test computational time and memory allocation for the standard scRNA-seq data analyses pipeline for one sample, including building metacells with SuperCell or with MetaCell followed by dimensionality reduction (PCA), clustering (Seurat) and DE analysis (t-test), scRNA-seq datasets of a different number of cells were generated by subsampling a larger dataset (GSE136831) [65]. DE analysis was performed for 3 clusters and 10′000 genes. The benchmarking was performed using the *time* linux command on an HPC cluster using 20 CPUs to allow parallel computation for MetaCell. To test the limits reached on standard desktop, the benchmarking was performed on an Intel core i7 with 16G of RAM.

### Building metacells that are consistent with cell type annotation

To build metacells that are consistent with a cell type annotation or specific conditions (e.g., treatments, donors) provided by the users, metacells that contain cells from different cell types or conditions are split such that each metacell contains cells from only one cell type and/or one condition.

## Declarations

### Ethics approval and consent to participate

Mouse experiments were performed following protocols approved by the Veterinary Authorities of the Canton Vaud according to Swiss law (animal license VD3612).

### Consent for publication

Not applicable.

### Availability of data and materials

The code to build and analyze metacells is available as *SuperCell* R package (https://github.com/GfellerLab/SuperCell).

Datasets analyzed in this study (Supplementary Table 1) are available in GEO under accession numbers GSE118767 (scRNA-seq of *cell_lines* [32]), GSE127465 (scRNA-seq of *TIICs* [4]), GSE116390 (scRNA-seq of *Cd8_TILs* [33]), GSE95315 (scRNA-seq of *brain_cells* [43]), GSM3852755 (scRNA-seq of *pancreatic_cells* [44]), GSE158055 (scRNA-seq of *COVID-19_atlas* [45]), GSE154763 (scRNA-seq of *TIM_atlas* [39], for the full list of integrated datasets, see Supplementary Table 2), GSE60424 (bulk RNA-seq of sorted CD4/CD8 T cells [62]), GSE86337 (bulk RNA-seq of 5 cancer cell lines [66]). ScRNA-seq data of purified T cells (*Tcells* [9]) are available from 10xGenomics [database](https://support.10xgenomics.com/single-cell-gene-expression/datasets). Bulk and scRNA-seq data of mouse data in treated and untreated conditions (*Mouse_DE* [35]) are available from Zenodo [database](https://doi.org/10.5281/zenodo.5048449) (zenodo.org/record/5048449).

### Authors’ contributions

DG designed the project. DG coordinated the project. MB developed the method. MB wrote the R package. MB, LT, AG, HM analyzed the data. CC performed the flow cytometry experiment. MJP supervised the flow cytometry experiments. MB and DG wrote the manuscript. SJC provided feedback on the method and the manuscript.

## Acknowledgements

Not applicable.

