## Supplementary Material for "Metacells untangle large and complex single-cell transcriptome networks"

### Supplementary Figures

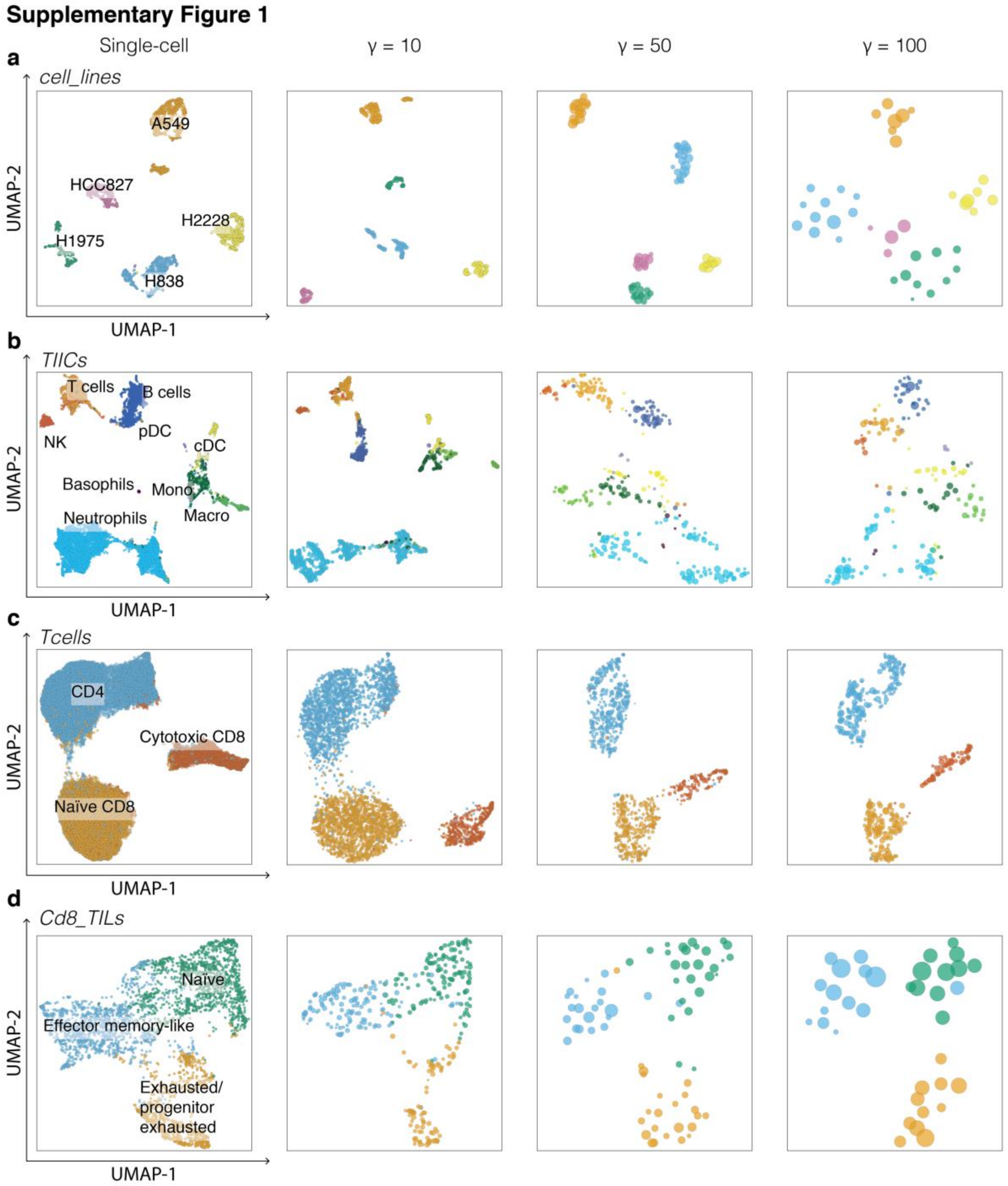

**Supplementary Figure 1. Metacells are compatible with UMAP visualization.** Examples of UMAPs of metacells at several graining levels. Colors indicate the initial cell type annotation and metacells are colored according to the majority of cells in each metacell. (**a**) Five cancer cell lines (*cell_lines*). (**b**) Tumor-infiltrating immune cells (*TIICs*). (**c**) T cells sorted from PBMC (*Tcells*). (**d**) Tumor-infiltrating CD8 T lymphocytes (*Cd8_TILs*).

**Supplementary Figure 2**

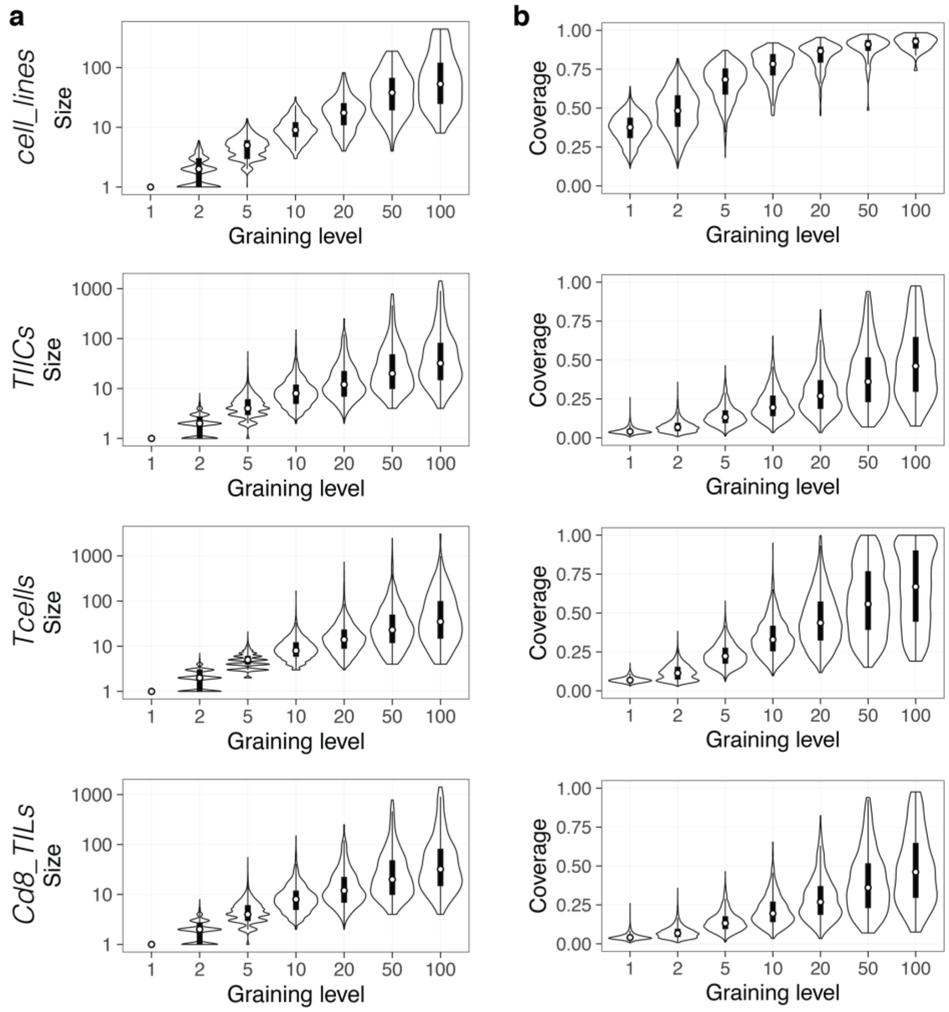

**Supplementary Figure 2. Metacell size and gene coverage distributions.**

**a**, Distribution of metacell sizes, defined as the number of single cells in each metacell. **b**, Distribution of metacell coverage, computed as the proportion of non-zero genes within each metacell. The white dots denote the median, and the boxes denote the interquartile range.

**Supplementary Figure 3**

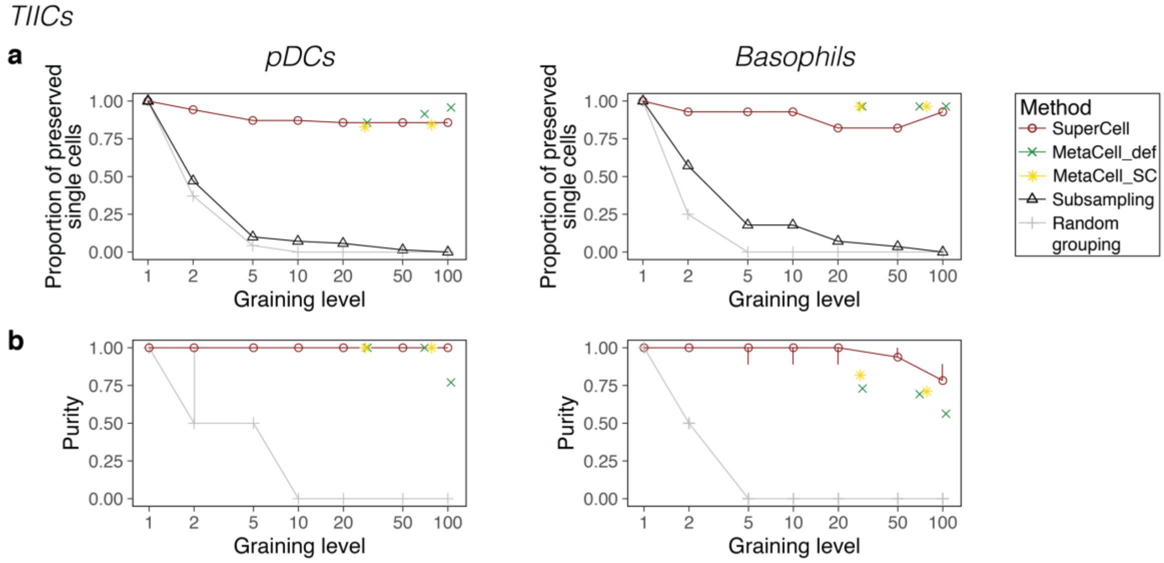

**Supplementary Figure 3. Metacells recover rare cell types.**

**a**, Proportion of correctly recovered single-cells from pDCs (left) and basophils (right), computed as a fraction of single cells of a particular cell types found in a metacell annotated to the same cell type. For the subsampling, it is computed as the proportion of subsampled single cells of the cell type of interest. **b**, Median purity of single cells within pDC metacells (left) or basophil metacells (right) (see Methods). Error bars denote the 1st and 3rd quartiles.

#### Supplementary Figure 4

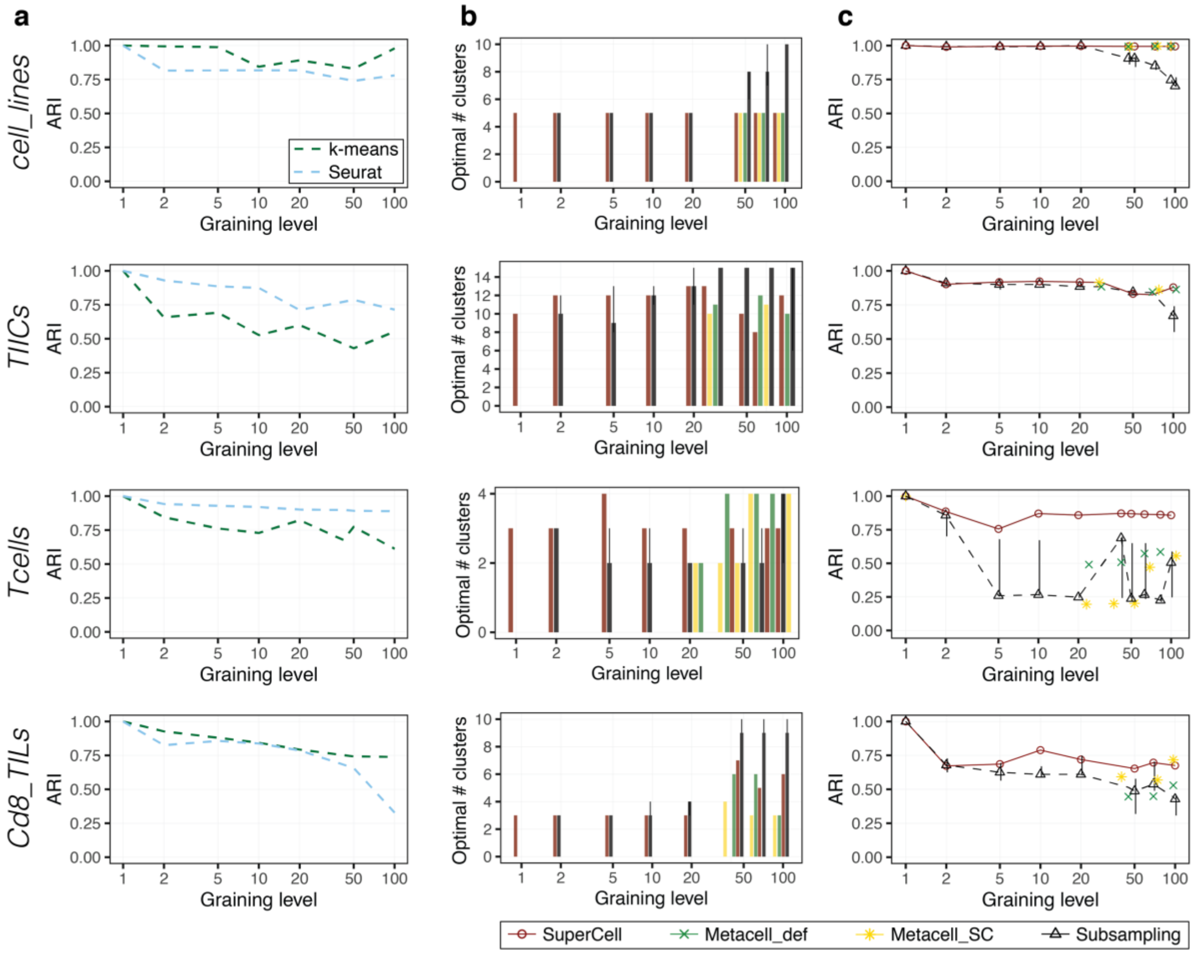

**Supplementary Figure 4. Metacells preserve clustering.**

**a**, Consistency of the metacell clustering obtained using k-means (green) and Seurat (blue) clustering algorithms. **b**, Optimal number of clusters based on the maximum silhouette coefficient. **c**, ARI values representing the consistency between clusters identified in metacells using the predicted optimal number of clusters (based on the maximum silhouette coefficient) and those identified in single cells.

**Supplementary Figure 5**

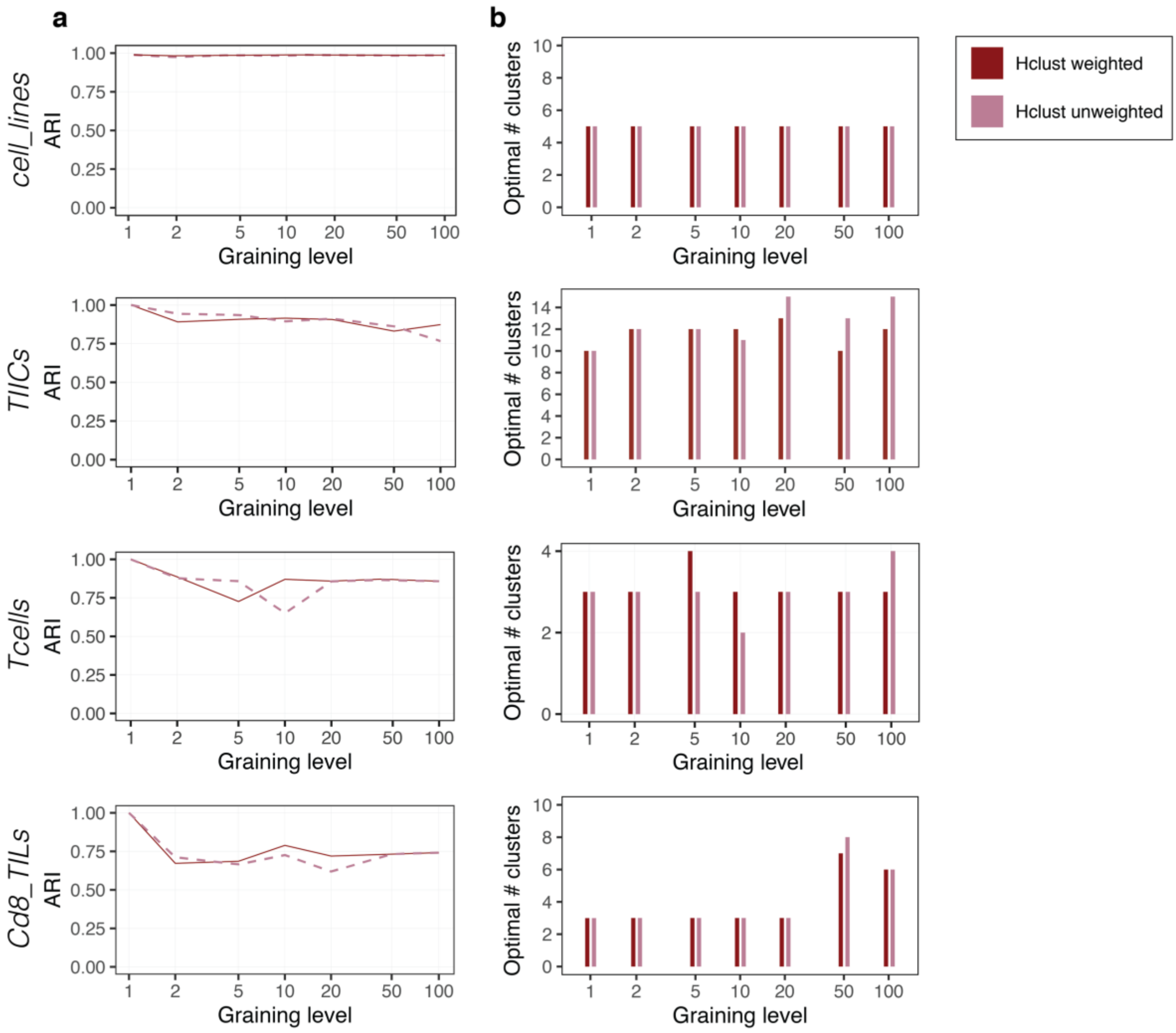

**Supplementary Figure 5. Metacells are compatible with unweighted clustering.** **a**, Consistency of the metacell clustering obtained using weighted (solid dark red line) and unweighted (dashed orchid line) hierarchical clustering algorithms. **b**, Optimal number of clusters based on the maximum silhouette coefficient for the weighted (dark red) and unweighted (orchid) hierarchical clustering.

#### Supplementary Figure 6

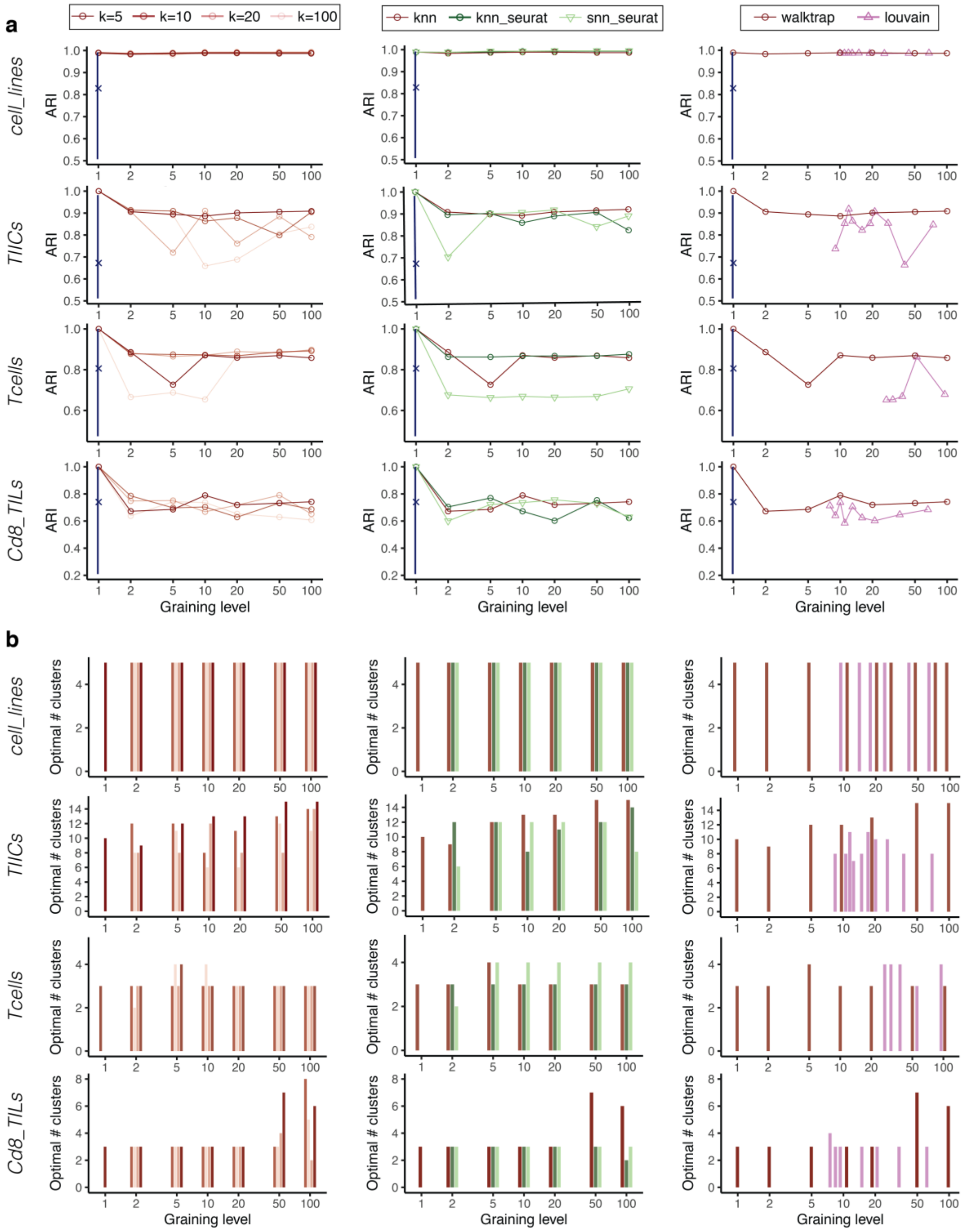

**Supplementary Figure 6. Clustering of metacells is robust to different ways of building metacells.**

Consistency of clustering (**a**) and optimal number of clusters (**b**) of metacells computed with different values for the parameter $k$ in kNN single-cell network (left), or different ways of the construction of single-cell network including Seurat kNN and Seurat sNN (shared nearest neighbors) (middle), or different ways of single-cell network clustering into metacells including Louvain algorithm for four datasets (right). The default parameters of the SuperCell algorithm are shown in dark red (i.e., kNN with $k = 5$ and walktrap clustering). The blue line shows the range of ARI values when other clustering algorithms are applied to the single-cell data (median shown with “X”).

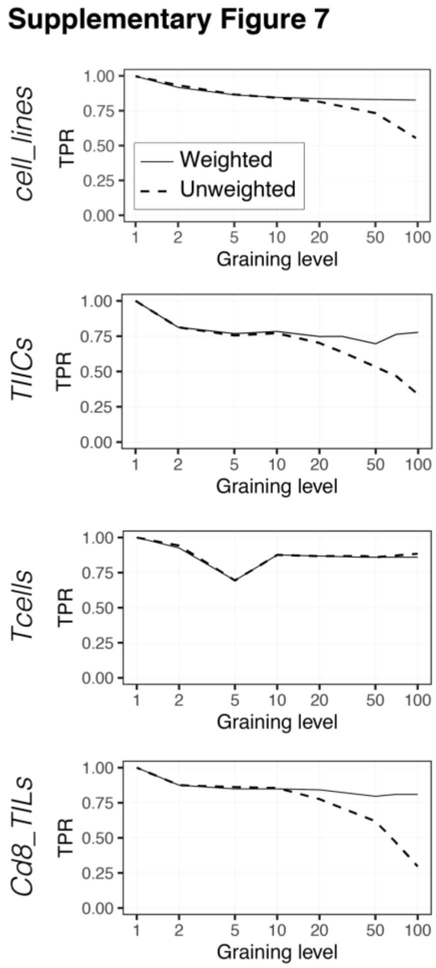

**Supplementary Figure 7. Metacells are compatible with unweighted differential expression analysis.**

Recovery of the cluster-specific differentially expressed genes using unweighted differential expression algorithm (i.e., unweighted t-test) for four datasets.

Supplementary Figure 8

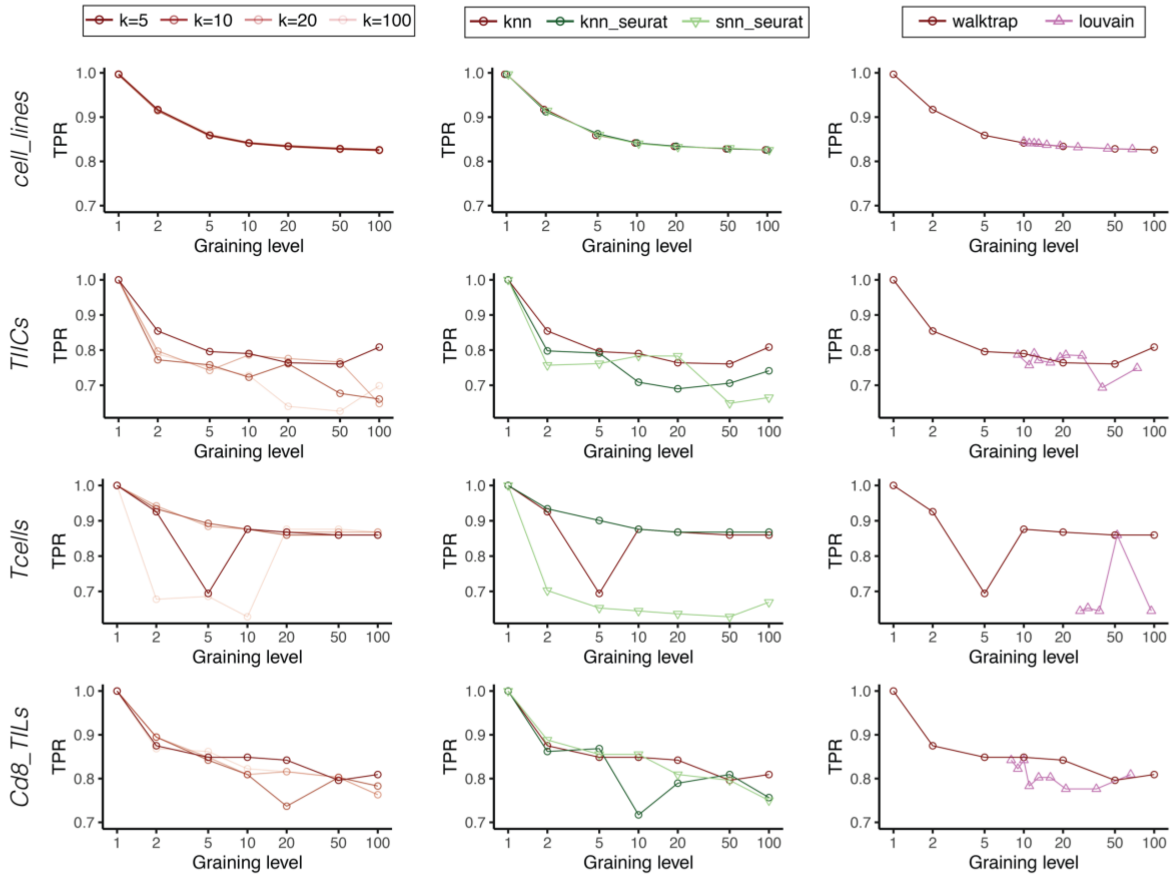

**Supplementary Figure 8. Differential expression of metacells is robust to different ways of building metacells.**

Recovery of cluster-specific differentially expressed genes of metacells computed with different values for the parameter $k$ in kNN single-cell network (left), or different ways of the construction of single-cell network, including Seurat kNN and Seurat sNN (shared nearest neighbors) (middle), or different ways of single-cell network clustering into metacells including Louvain algorithm for four datasets (right). The default parameters of the SuperCell algorithm are shown in dark red (i.e., kNN with $k = 5$ and walktrap clustering).

#### Supplementary Figure 9

*Mouse_DE*

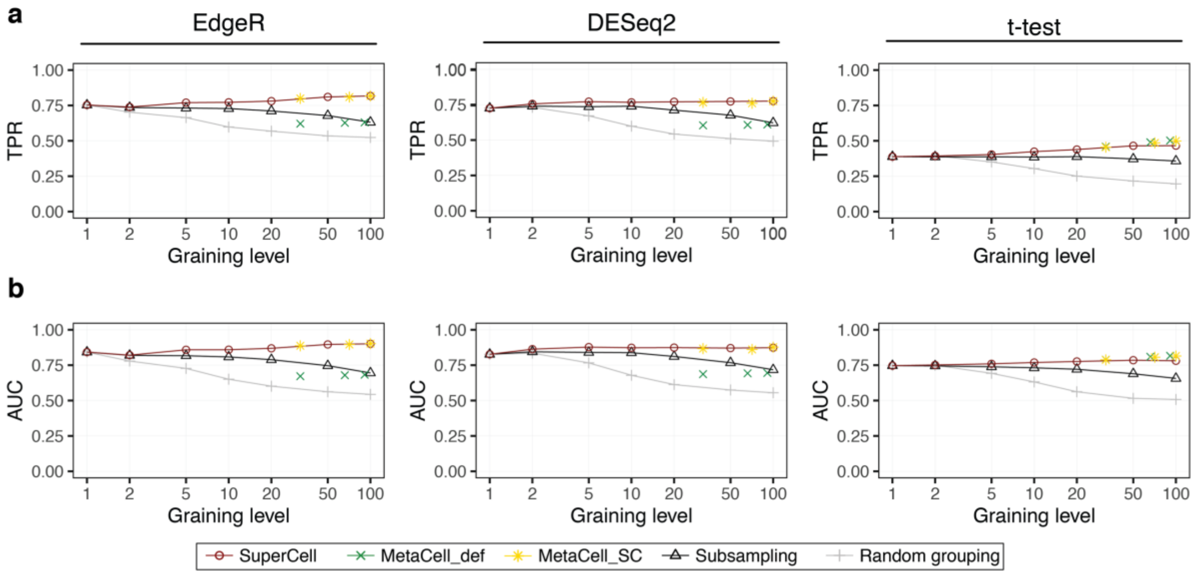

**Supplementary Figure 9. Metacells preserve differential expression between conditions.**

TPR (**a**) and AUC (**b**) of the recovery of differentially expressed genes between treated and control samples using EdgeR, DESeq2 and t-test approaches. The ground truth is the differential expression analysis from bulk RNA-seq.

**Supplementary Figure 10**

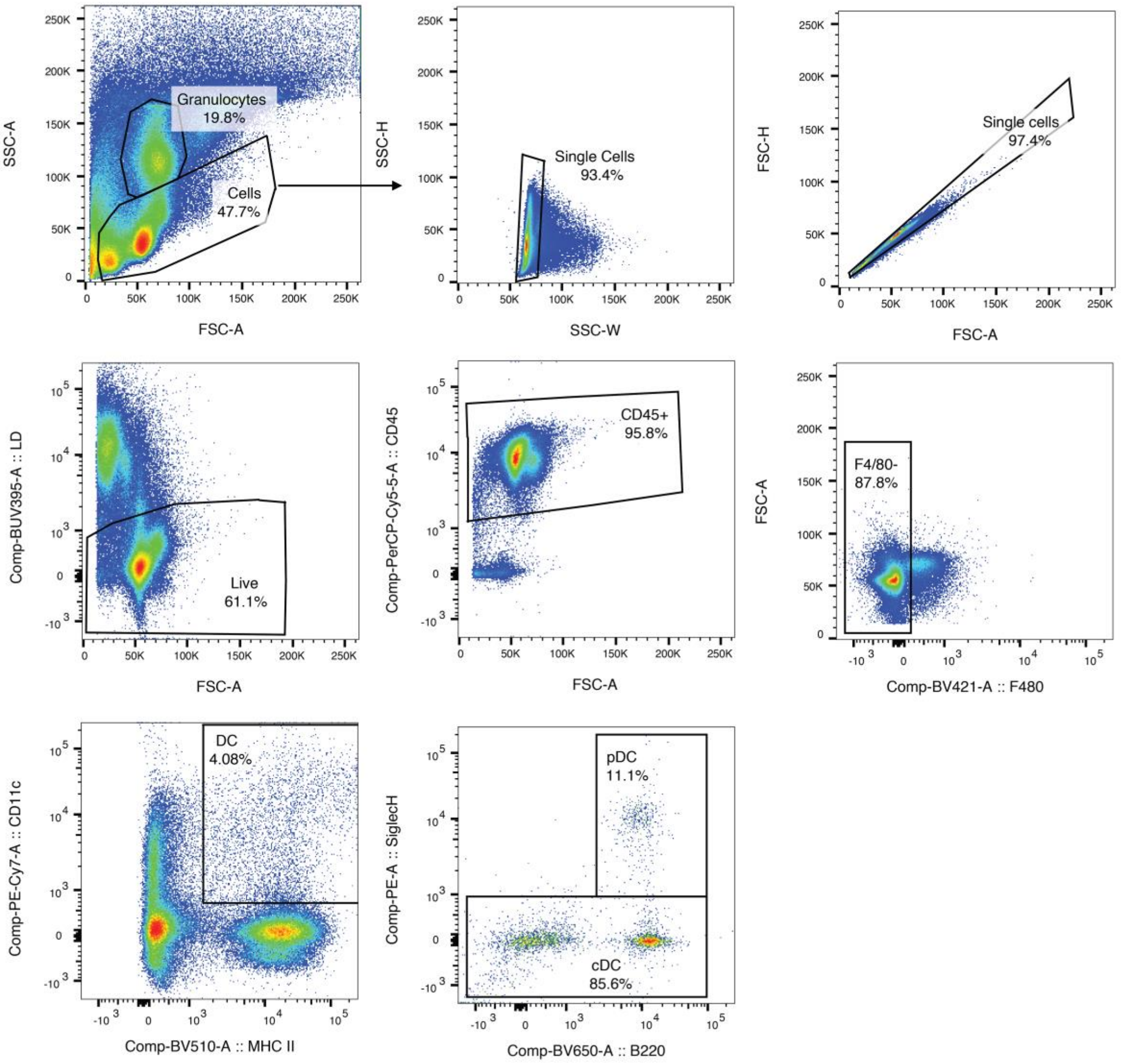

**Supplementary Figure 10. Gating strategy for the flow cytometry analysis of DCs from murine KP1.9 lung adenocarcinoma for one representative sample.**

**Supplementary Figure 11**

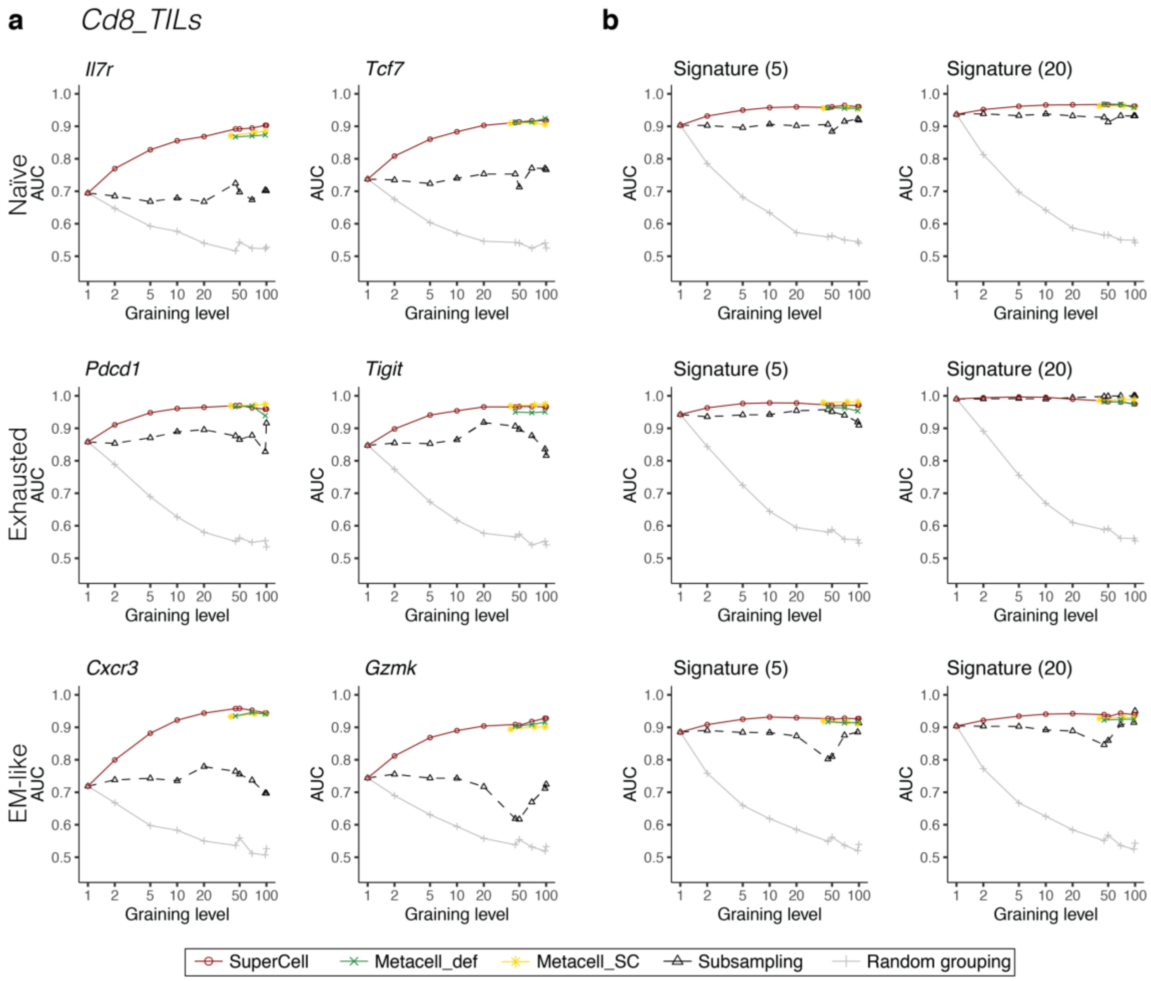

**Supplementary Figure 11. Cell type annotation in the *Cd8_TILs* dataset.**

**a**-**b**, AUC of the recovery of naïve (top), exhausted/progenitor exhausted (middle) and effector memory-like (bottom) CD8 T cells from the *TILs* dataset using single markers (**a**) or signatures defined from the same dataset (**b**) that consists of the top 5 or 20 upregulated genes.

**Supplementary Figure 12**

**a** *brain_cells*

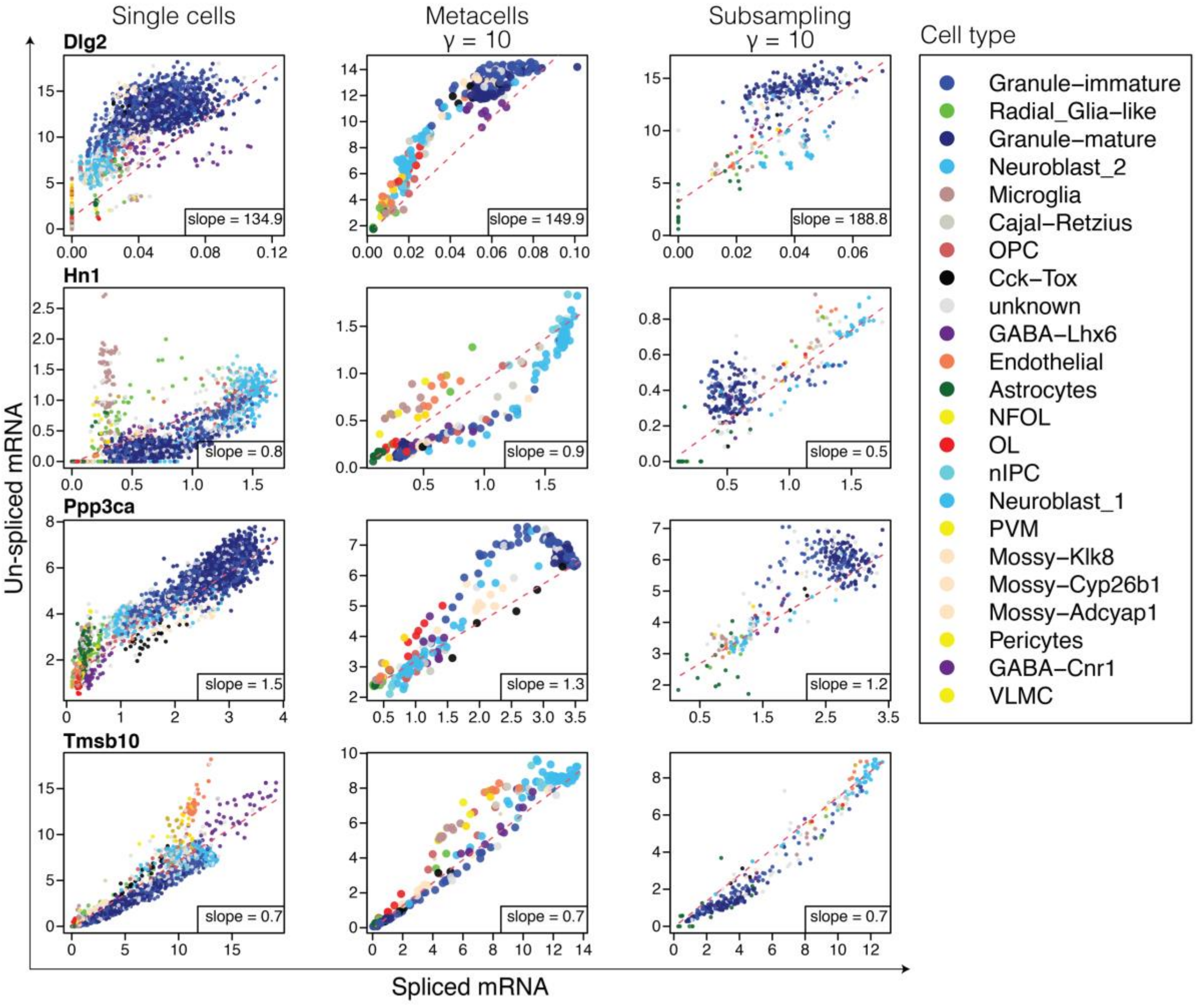

**b**

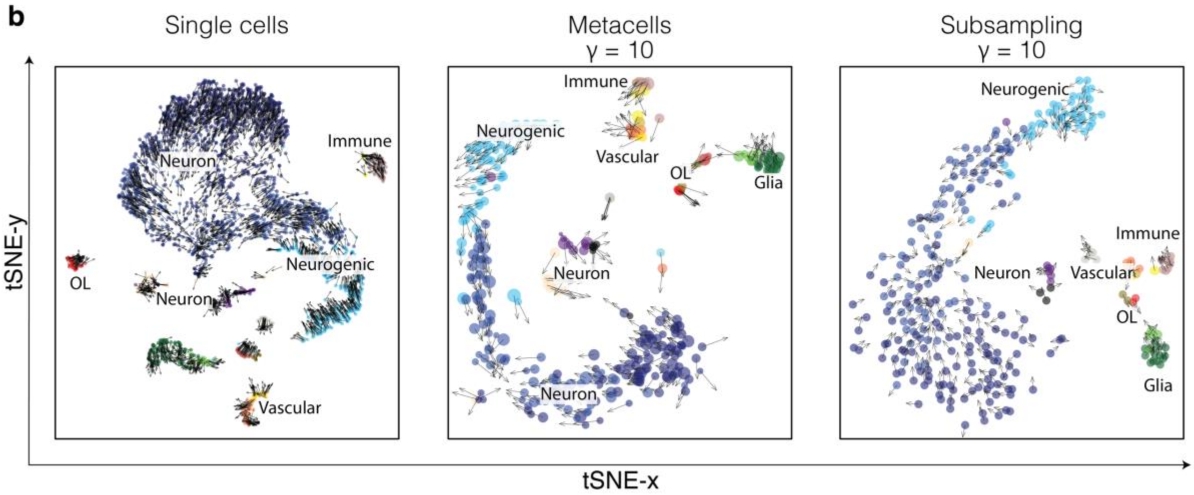

**Supplementary Figure 12. Conservation of RNA velocity results in the *brain_cells* dataset.**

**a**, Spliced/un-spliced phase portraits and estimated equilibrium slopes (red dashed lines) for the single cells (left), metacells ($\gamma = 10$) (middle) and subsampling ($\gamma = 10$) (right) in the *brain_cell* dataset. **b**, Separate tSNE visualization of RNA velocity for the single cells (left), metacells ($\gamma = 10$) (middle) and subsampling ($\gamma = 10$) (right). Colors indicate the cell type annotation of single-cell data and metacells are colored according to the majority of cells in each metacell.

#### Supplementary Figure 13

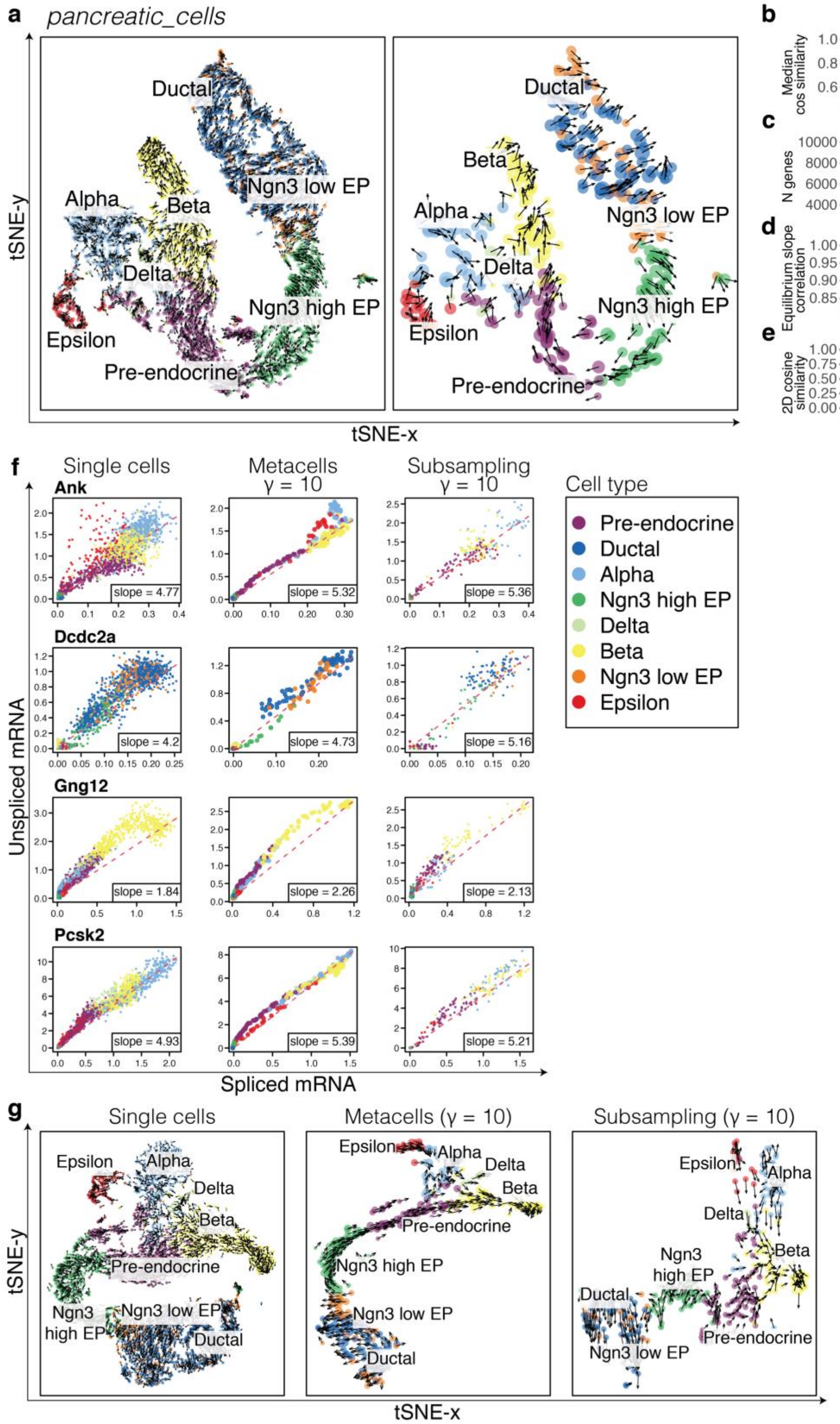

**Supplementary Figure 13. Conservation of RNA velocity results in the *pancreatic_cells* dataset.**

**a**, Joint tSNE visualization of RNA velocity for single cells (left) and metacells ($\gamma = 10$) (right) in the *pancreatic_cells* dataset ($N = 3'696$). Colors indicate the cell type annotation of single-cell data and metacells are colored according to the majority of cells in each metacell. **b**, Median purity of metacell velocity computed as a cosine similarity of velocities within each metacell. **c**, Number of genes with valid estimated equilibrium slope values. **d**, Median Pearson correlation of gene equilibrium slope values obtained in single cells and metacells. **e**, Median similarity of 2D RNA velocities computed as a cosine similarity between a 2D RNA velocity of each single cell and a 2D RNA velocity of the metacell it belongs to. **f**, Spliced/un-spliced phase portraits and estimated equilibrium slopes (red dashed lines) for the single cells (left), metacells ($\gamma = 10$) (middle) and subsampling ($\gamma = 10$) (right). **g**, Separate tSNE visualization of RNA velocity for the single cells (left), metacells ($\gamma = 10$) (middle) and subsampling ($\gamma = 10$) (right).

**Supplementary Figure 14**

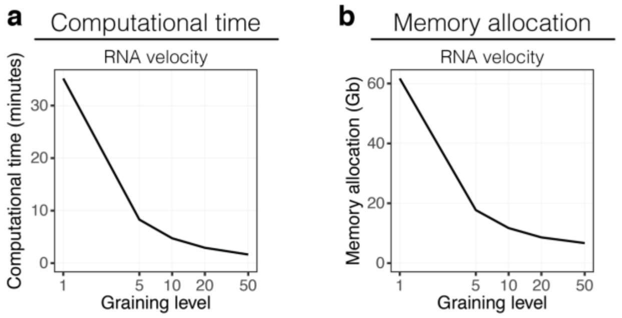

**Supplementary Figure 14. Computational time and memory allocation for RNA velocity.**

**a**-**b**, Computational time (**a**) and memory allocation (**b**) for the RNA velocity over different graining levels for the *brain_cells* dataset ($N = 3'396$).

#### Supplementary Figure 15

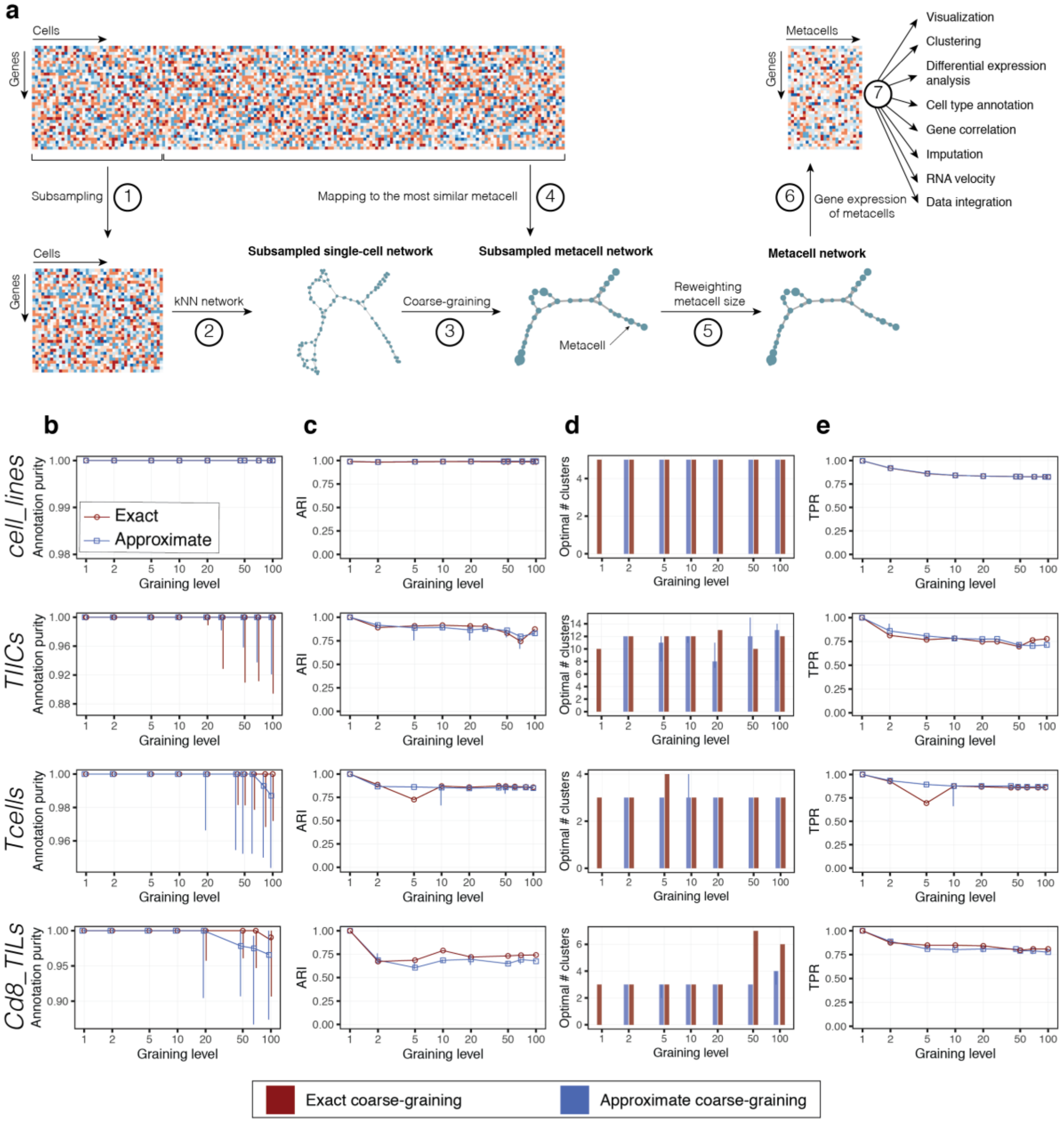

**Supplementary Figure 15. Approximate coarse-graining in SuperCell.**

**a**, Overview of the approximate coarse-graining approach. (1) A subset of cells is randomly selected. (2) The single-cell network is constructed from the subsampled single-cell gene expression matrix using k-nearest neighbors (kNN) algorithm. (3) The metacell network is constructed by grouping similar cells into metacells at a user-defined graining level ($\gamma$). (4) Remaining cells are mapped to the most similar metacell. (5) The metacell network is reweighted. (6) The gene expression matrix of metacells is computed by averaging gene expression within each metacell. (7) This metacell gene expression matrix can be used for visualization and downstream analyses such as clustering, differential expression, cell type annotation, gene correlation, RNA velocity and data integration. **b-d**, Comparison of the results of the exact and the approximate coarse-graining in terms of metacell purity (**b**), consistency of clustering (**c**), optimal number of clusters (**d**) and recovery of cluster-specific differentially expressed genes (**e**) for the exact (red) and the approximate (blue) coarse-graining. For the approximate coarse-graining, the center of the error bars denotes the median, and the extrema denote the $1^{st}$ and $3^{rd}$ quartiles (obtained with different random seeds).

#### Supplementary Figure 16

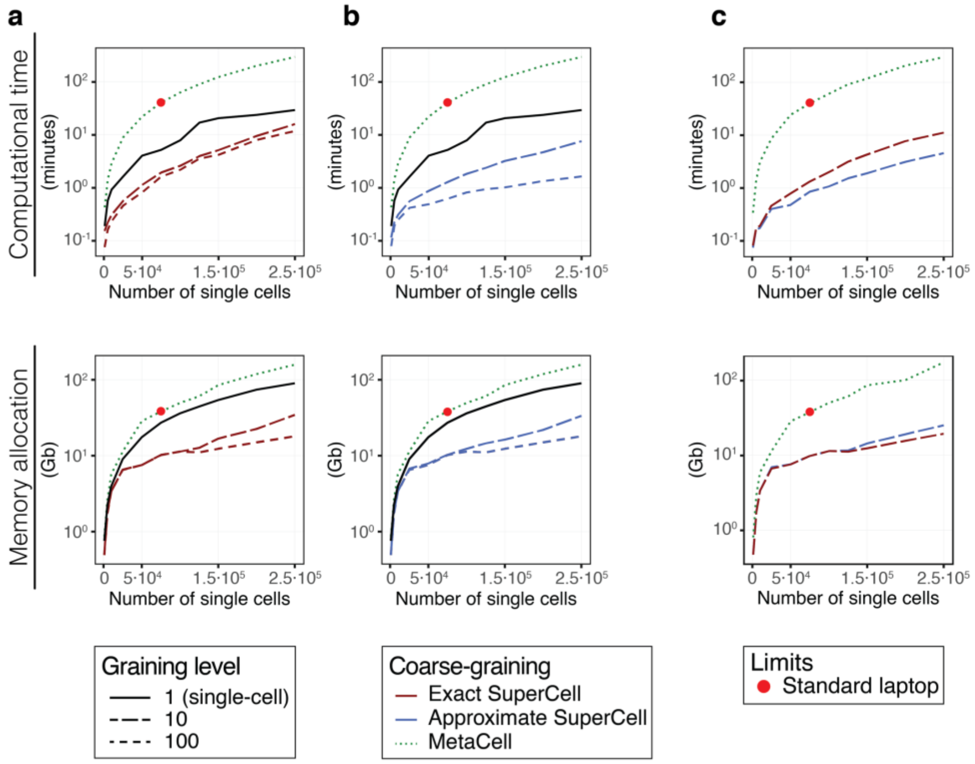

**Supplementary Figure 16. Computational time and memory allocation for metacell construction and downstream analyses.**

**a**,**b**, Computational time (top) and memory allocation (bottom) for the building of metacells with MetaCell, exact (**a**) or the approximate (**b**) SuperCell followed by downstream analyses including dimensionality reduction, clustering and DE analysis for the metacell and the single-cell data on a single dataset (extracted from GSE136831, see Methods). **c**, Computational time (top) and memory allocation (bottom) for the building of metacells with MetaCell or the exact and approximate SuperCell. Red dots represent the limits reached on standard desktops (16G of RAM).

#### Supplementary Table 1. Datasets used for the analysis.

**Single-cell RNA-seq dataset**

| Dataset_ID | Cell type | Organism | Number of cells | Pipeline demonstration | Ground truth of cell type available | Gene-gene correaltion | RNA Velocity | PMID | GEO |
|---|---|---|---|---|---|---|---|---|---|
| *cell_lines* | A549/H838/H1975/H2228/HCC827 | human | 3'918 | x | x | x | | 31133762 | GSE118767 |
| *TIICs* | pDC/cDC/Macrophafges/Monocytes/Neutrophils/Tcells/Bcells | mouse | 15'939 | | | | | 30979687 | GSE127465 |
| *Tcells* | CD4/CD8 T cells | human | 40'560 | x | x | | | 28091601 | 10x Datasets |
| *Cd8_TILs* | CD8 T cells | mouse | 3'574 | x | | | | 32313720 | GSE116390 |
| *Mouse_DE* | treated/untreated | mouse | 16'882 | | x | | | 34584091 | Zenodo dataset |
| *brain_cells* | Neuron/Glia/Vascular/Immune | mouse | 3'396 | | | | x | 29335606 | GSE95315 |
| *pacreatic_cells* | endocrine cells from embryonic day 15.5 | mouse | 3'696 | | | | x | 31160421 | GSM3852755 |
| *COVID-19_atlas* | Tcells/NK/Bcells/Myeloid/Epithelial | human | 1'462'702 | | | | | 33657410 | GSE158055 |
| *TIM_atlas* | pDC/cDC/Macrophafges/Monocytes | human | 108'566 | | | | | 33545035 | GSE154763 |

**Bulk RNA-seq dataset**

| Dataset_ID | Cell type | Organism | Number of samples | | | | | PMID | GEO |
|---|---|---|---|---|---|---|---|---|---|
| *cell_lines_bulk* | A549/H838/H1975/H2228/HCC827 | human | 10 | | | | | 27899618 | GSE86337 |
| *Tcells_bulk* | CD4/CD8 T cells | human | 40 | | | | | 25314013 | GSE60423 |
| *Mouse_DE_bulk* | treared/untreated | mouse | 6 | | | | | 34584091 | Zenodo dataset |

#### Supplementary Table 2. Datasets integrated in the *TIM_atlas* dataset.

| Protocol | Study | PMID | Cancer |
|---|---|---|---|
| 10x 5' | Cheng et al., 2021 | 33545035 | Esophageal carcinoma (ESCA) |
| | | | ovarian or fallopian tube carcinoma (OV-FTC) |
| | | | Pancreatic adenocarcinoma (PAAD) |
| | | | Thyroid carcinoma (THCA) |
| | | | Uterine Corpus Endometrial Carcinoma (UCEC) |
| | | | Lymphoma (LYM) |
| | | | Myeloma (MYE) |
| | | | Renal cancer (KIDNEY) |
| 10x 3' | Lambrechts et al., 2018 | 29988129 | Lung cancer (LUNG) |
| | Peng et al., 2019 | 31273297 | Pancreatic adenocarcinoma (PAAD) |
| | Zhang et al., 2020 | 32302573 | Colorectal cancer (CRC) |
| inDrop | Zilionis et al., 2019 | 30979687 | Lung cancer (LUNG) |
| | Azizi et al., 2018 | 29961579 | Breast cancer (BRCA) |
| Smart-seq2 | Zhang et al., 2020 | 32302573 | Colorectal cancer (CRC) |

Datasets for integration and integration approach taken from Cheng et al., 2021

#### Supplementary Table 3. Genes ranked better in differential expression analysis (cDC vs pDC) at metacell level.

| Gene | p.value | adj.p.value | pct.1 | pct.2 | logFC | w.mean.1 | w.mean.2 | rank at single-cell | rank at super-cell | delta | Cell membrane | Available antibody |
|---|---|---|---|---|---|---|---|---|---|---|---|---|
| Actg1 | 0 | 0 | 1 | 1 | 1.605329 | 3.007947 | 1.456752 | 9 | 4 | -5 | | |
| Tmsb4x | 0 | 0 | 1 | 1 | 1.555013 | 3.9163 | 2.38875 | 8 | 5 | -3 | | |
| **H2-Aa** | **0** | **0** | **1** | **1** | **1.535628** | **2.833846** | **1.497637** | **100** | **6** | **-94** | **x** | **x** |
| Crip1 | 0 | 0 | 1 | 1 | 1.450344 | 1.68243 | 0.294961 | 12 | 7 | -5 | | |
| **Cd74** | **0** | **0** | **1** | **1** | **1.239884** | **2.832465** | **1.684596** | **96** | **12** | **-84** | **x** | **x** |
| Ifi30 | 0 | 0 | 1 | 1 | 1.00315 | 1.376976 | 0.438663 | 92 | 17 | -75 | | |
| Pim1 | 0 | 0 | 1 | 1 | 0.881269 | 1.226782 | 0.3604 | 114 | 20 | -94 | x | |
| Actb | 0 | 0 | 1 | 1 | 0.81907 | 3.36918 | 2.62061 | 155 | 23 | -132 | x | x |
| Calm1 | 0 | 0 | 1 | 1 | 0.794634 | 1.267894 | 0.593374 | 124 | 24 | -100 | x | x |
| Ftl1 | 0 | 0 | 1 | 1 | 0.775703 | 1.932204 | 1.18621 | 120 | 27 | -93 | | |
| Gm10116 | 0 | 0 | 1 | 1 | 0.741254 | 1.848595 | 1.135099 | 123 | 29 | -94 | | |
| Vim | 0 | 0 | 1 | 1 | 0.730312 | 0.895743 | 0.233075 | 202 | 30 | -172 | x | x |
| S100a11 | 0 | 0 | 1 | 1 | 0.663528 | 0.96415 | 0.305091 | 83 | 33 | -50 | | |
| H2afz | 0 | 0 | 1 | 1 | 0.642373 | 0.868661 | 0.28277 | 98 | 35 | -63 | | |
| Sh3bgrl3 | 0 | 0 | 1 | 1 | 0.640981 | 1.374543 | 0.733037 | 130 | 36 | -94 | | |
| S100a6 | 0 | 0 | 1 | 1 | 0.640428 | 1.216493 | 0.764779 | 228 | 37 | -191 | x | x |
| Tmsb10 | 0 | 0 | 1 | 1 | 0.636512 | 1.122315 | 0.547541 | 160 | 38 | -122 | | |
| Cd52 | 0 | 0 | 1 | 1 | 0.619723 | 1.083616 | 0.579177 | 182 | 40 | -142 | x | |
| Slc45a3 | 0 | 0 | 1 | 1 | 0.597725 | 0.782665 | 0.228264 | 99 | 42 | -57 | | |
| Myl6 | 0 | 0 | 1 | 1 | 0.587303 | 1.220488 | 0.652583 | 154 | 43 | -111 | | |
| Pfn1 | 0 | 0 | 1 | 1 | 0.586644 | 1.276556 | 0.772013 | 231 | 44 | -187 | | |
| Zfp36 | 0 | 0 | 1 | 1 | 0.557965 | 1.327452 | 0.817545 | 277 | 51 | -226 | | |
| Lrrc58 | 0 | 0 | 1 | 1 | 0.531825 | 1.107848 | 0.646132 | 227 | 55 | -172 | | |
| Junb | 0 | 0 | 1 | 1 | 0.51861 | 0.767571 | 0.278914 | 181 | 57 | -124 | | |
| Tubb5 | 0 | 0 | 1 | 1 | 0.506081 | 0.888655 | 0.428209 | 187 | 59 | -128 | | |
| Cytip | 0 | 0 | 1 | 1 | 0.495787 | 0.963034 | 0.543827 | 248 | 60 | -188 | | |
| Arpc2 | 0 | 0 | 1 | 1 | 0.419255 | 0.950488 | 0.547355 | 233 | 76 | -157 | x | |
| Tuba1b | 0 | 0 | 1 | 1 | 0.400753 | 0.72402 | 0.362144 | 205 | 80 | -125 | | |
| Pcbp2 | 0 | 0 | 1 | 1 | 0.398435 | 0.937415 | 0.589119 | 240 | 81 | -159 | | |
| Anxa5 | 0 | 0 | 1 | 1 | 0.380112 | 0.516261 | 0.151547 | 219 | 88 | -131 | x | x |
| Spi1 | 0 | 0 | 1 | 1 | 0.35251 | 0.472239 | 0.132607 | 173 | 93 | -80 | | |
| Gm4750 | 0 | 0 | 1 | 1 | 0.343985 | 0.650829 | 0.318748 | 177 | 99 | -78 | | |
| Gm8464 | 0 | 0 | 1 | 1 | 0.343985 | 0.650829 | 0.318748 | 178 | 100 | -78 | | |
| Iscu | 0 | 0 | 1 | 1 | 0.339716 | 0.42957 | 0.108064 | 109 | 102 | -7 | | |
| Samhd1 | 0 | 0 | 1 | 1 | 0.33855 | 0.570063 | 0.24053 | 207 | 105 | -102 | x | x |
| Psme2 | 0 | 0 | 1 | 1 | 0.337769 | 0.514583 | 0.199942 | 183 | 106 | -77 | | |
| Cfl1 | 0 | 0 | 1 | 1 | 0.313879 | 1.542907 | 1.278436 | 271 | 112 | -159 | x | x |
| Clic4 | 2.22E-16 | 6.18E-12 | 1 | 1 | 0.305731 | 0.482597 | 0.195308 | 200 | 150 | -50 | x | x |

Experimentally tested genes

#### Supplementary Table 4. Genes ranked better in differential expression analysis (pDC vs cDC) at metacell level.

| Gene | p.value | adj.p.value | pct.1 | pct.2 | logFC | w.mean.1 | w.mean.2 | rank at single-cell | rank at super-cell | delta | Cell membrane | Available antibody |
|---|---|---|---|---|---|---|---|---|---|---|---|---|
| **Ly6e** | **0** | **0** | **1** | **0.953** | **1.480381** | **2.002309** | **0.480345** | **7** | **4.00** | **-3** | **x** | **x** |
| Irf8 | 0 | 0 | 1 | 1 | 1.468721 | 2.13397 | 0.518266 | 8 | 5.00 | -3 | | |
| Serinc3 | 0 | 0 | 1 | 0.97 | 1.156719 | 1.449309 | 0.331378 | 21 | 14.00 | -7 | x | |
| Gnas | 0 | 0 | 1 | 1 | 0.791572 | 1.555245 | 0.835275 | 58 | 25.00 | -33 | x | |
| Unc93b1 | 0 | 0 | 1 | 0.981 | 0.62 | 1.014853 | 0.373748 | 44 | 34.00 | -10 | | |
| Dnajc7 | 0 | 0 | 1 | 1 | 0.619044 | 0.746032 | 0.132681 | 40 | 36.00 | -4 | | |
| Irf2bp2 | 0 | 0 | 1 | 0.981 | 0.534315 | 0.769351 | 0.25251 | 49 | 46.00 | -3 | | |
| Hsp90b1 | 0 | 0 | 1 | 1 | 0.532243 | 0.745137 | 0.215259 | 52 | 49.00 | -3 | | |
| Ly86 | 0 | 0 | 1 | 1 | 0.523536 | 0.862894 | 0.364407 | 87 | 50.00 | -37 | | |
| Smim14 | 0 | 0 | 1 | 0.979 | 0.50654 | 0.692369 | 0.200778 | 99 | 52.00 | -47 | | |
| **Cd47** | **0** | **0** | **1** | **0.989** | **0.50137** | **1.19715** | **0.678006** | **66** | **53.00** | **-13** | **x** | **x** |
| Tagln2 | 0 | 0 | 1 | 0.989 | 0.415992 | 1.366392 | 0.909724 | 152 | 68.00 | -84 | | |
| Tmed2 | 0 | 0 | 1 | 0.985 | 0.412263 | 0.663359 | 0.269782 | 100 | 69.00 | -31 | | |
| Ppia | 0 | 0 | 1 | 1 | 0.403637 | 2.224881 | 1.81784 | 1000 | 70.00 | -930 | | |
| Laptm5 | 0 | 0 | 1 | 1 | 0.400209 | 1.407769 | 0.951309 | 287 | 72.00 | -215 | | |
| Krtcap2 | 0 | 0 | 1 | 0.97 | 0.389737 | 0.599857 | 0.215232 | 103 | 75.00 | -28 | x | |
| Rnaset2b | 0 | 0 | 1 | 0.974 | 0.389239 | 0.897053 | 0.491162 | 241 | 76.00 | -165 | | |
| Ubc | 0 | 0 | 1 | 1 | 0.347795 | 1.032611 | 0.659598 | 1000 | 88.00 | -912 | | |
| Rnaset2a | 0 | 0 | 1 | 0.974 | 0.346367 | 0.914968 | 0.542602 | 299 | 89.00 | -210 | | |
| **Cd44** | **3.3E-15** | **9.2703E-11** | **1** | **1** | **0.363489** | **0.665219** | **0.310748** | **246** | **142.00** | **-104** | **x** | **x** |

Experimentally tested genes

**Supplementary Table 5. Antibodies used in flow cytometry (Fig. 2f, Supplementary Fig. 11).**

| Marker | Fluorochrome | Clone | Dilution | Cat | Brand |
|---|---|---|---|---|---|
| CD45 | PercP-Cy5.5 | 30-F11 | 200 | 45-0451-82 | eBioscience |
| CD44 | BV605 | IM7 | 150 | 103047 | BioLegend |
| F4/80 | BV421 | BM8 | 100 | 123132 | BioLegend |
| CD11c | Pe-Cy7 | N418 | 200 | 117318 | BioLegend |
| MHC II (I-A/I-E) | BV510 | M5/114/15/2 | 200 | 107636 | BioLegend |
| Ly6A/E | BV711 | D7 | 150 | 108131 | BioLegend |
| B220 | BV650 | RA3-6B2 | 150 | 103241 | BioLegend |
| SiglecH | PE | 551 | 150 | 129605 | BioLegend |
| CD47 | APC-Cy7 | miap301 | 150 | 127525 | BioLegend |
| CD74 | AF647 | In1/CD74 | 150 | 151003 | BioLegend |
